# p53-mediated neurodegeneration in the absence of the nuclear protein Akirin2

**DOI:** 10.1101/2021.07.05.451155

**Authors:** Stacey L. Peek, Peter J. Bosch, Ethan Bahl, Brianna J. Iverson, Mrutyunjaya Parida, Preeti Bais, J. Robert Manak, Jacob J. Michaelson, Robert W. Burgess, Joshua A. Weiner

## Abstract

Proper gene regulation is critical for both neuronal development and maintenance as the brain matures. We previously demonstrated that Akirin2, an essential nuclear protein that interacts with transcription factors and chromatin remodeling complexes, is required for the embryonic formation of the cerebral cortex. Here we show that Akirin2 plays a mechanistically distinct role in maintaining healthy neurons during cortical maturation. Restricting Akirin2 loss to excitatory cortical neurons resulted in progressive neurodegeneration via necroptosis and severe cortical atrophy with age. Comparing transcriptomes from Akirin2-null postnatal neurons and cortical progenitors revealed that targets of the tumor suppressor p53, a regulator of both proliferation and cell death encoded by *Trp53*, were consistently upregulated. Heterozygous deletion of *Trp53* rescued neurodegeneration in Akirin2-null neurons. These data: 1) implicate Akirin2 as a critical neuronal maintenance protein; 2) identify p53 pathways as mediators of Akirin2 functions; and 3) suggest Akirin2 dysfunction may be relevant to neurodegenerative diseases.

## Introduction

During development, neural progenitor cells become mitotically quiescent and terminally differentiate into a multitude of specific neuronal subtypes. Maintaining subtype identity and postmitotic status requires continuous expression of the proper gene patterns throughout a neuron’s long postmitotic life (Deneris et al., 2014; Cholewa-Waclaw et al., 2016; Gallegos et al., 2018). This includes both activating neuronal cell-type specific genes as well as suppressing non-neuronal and cell cycle-related genes. Dysregulation of gene networks in postmitotic neurons can lead to defects in innervation (Lin et al., 1998; Arber et al., 2000; Kania et al., 2003; Chen et al., 2013), dendritic and axonal deterioration (Kadkhodaei et al., 2013; Lipinski et al., 2020), loss of cell type identity (Liu et al., 2010; Bovetti et al., 2013; Montana et al., 2013), neuronal network dysfunction (Chen et al., 2013; Kadkhodaei et al., 2013; Lipinski et al., 2020) and cell death (Ninkovic et al., 2010; von Schimmelmann et al., 2016). Thus, it is not surprising that disrupted gene regulation is associated with memory impairment (Barrett et al., 2011; Vogel-Ciernia et al., 2013), cognitive dysfunction, and neurodegenerative diseases (De Jager et al., 2014; Sanchez-Mut et al., 2016; Watson et al., 2016; Berson et al., 2018; Li et al., 2019). In fact, many studies of neurodegenerative diseases find dysregulation of gene expression in postmitotic neurons prior to symptom onset, implicating gene dysregulation as the earliest neuronal insult in these diseases (Yang et al., 2003; De Jager et al., 2014). Cell cycle genes in particular are ectopically expressed in neurodegenerative diseases, and postmitotic neurons gain phenotypes suggestive of re-entry into the cell cycle (Busser et al., 1998; Yang et al., 2003; McShea et al., 2007; Lee et al., 2009; Wang et al., 2009; Li et al., 2019). As postmitotic neurons are physically incapable of dividing and completing the cell cycle, this re-entry typically leads to cell death (Park et al., 2007). Proper regulation of gene expression patterns is critical for preventing such aberrant cell cycle re-entry and thus for postmitotic neuron maintenance and survival.

Akirin2 (Aki2) is a highly-conserved, nuclear protein that regulates gene expression by interacting with transcription factors (Nowak et al., 2012; Bonnay et al., 2014; Tartey et al., 2014) and members of chromatin remodeling complexes (Giot et al., 2003; Nowak et al., 2012; Bonnay et al., 2014; Tartey et al., 2014; Tartey et al., 2015; Liu et al., 2017); reviewed by (Bosch et al., 2020). Invertebrates such as *C. elegans* and *Drosophila* harbor a single Akirin gene in their genomes, whilst mammals have two homologues (*Akirin1* and *Akirin2*). Goto et al. (2008) generated both *Akirin1* (*Aki1*) and *Akirin2* (*Aki2*) null mice; whereas the former were viable and fertile with no outward defects, constitutive *Aki2* knockouts exhibited early embryonic lethality, identifying Aki2 as the critical mammalian homologue (Goto et al., 2008). Utilizing an *Emx1-Cre* driver to delete a conditional *Aki2* mutant allele in forebrain neural progenitor cells, we previously found an essential role for Aki2 in cerebral cortex development. Without Aki2, neural progenitor cells exited the cell cycle prematurely which was followed swiftly by massive apoptosis, resulting in few if any cortical neurons after embryonic day (E) 13. As a result of cortical agenesis, these mice were microcephalic and nearly all died shortly after birth (Bosch et al., 2016). This work identified previously-unknown roles for Akirins in embryonic brain development that are consistent with prior and subsequent studies indicating that Aki2 regulates proliferation and differentiation while inhibiting apoptosis in a variety of tissues and cell types (Komiya et al., 2008; Krossa et al., 2015; Tartey et al., 2015; Liu et al., 2017; Bosch et al., 2018; Bosch et al., 2019; Leng et al., 2019).

Here, we show that Aki2 expression continues throughout cortical development, albeit at lower levels in postnatal neurons than in the embryonic telencephalon. As postmitotic neurons have progressed beyond the stages of proliferation and differentiation, we hypothesized that residual Aki2 expression in the maturing postnatal brain indicates distinct roles in postmitotic neurons. To test this hypothesis, we utilized a well-characterized *CaMKIIα-Cre* driver (Tsien et al., 1996) to disrupt the *Aki2* gene in many postmitotic excitatory neurons of the cortex beginning around postnatal day (P)18. In contrast to embryonic cortical phenotypes in which massive apoptosis occurred within 24 hours of Cre expression (Bosch et al., 2016), we found that *Aki2* mutants are not immediately affected. Rather, we present evidence that they undergo a slow process of death by necroptosis over several months in a pattern that suggests initial death is cell-autonomous and that neighboring Cre-negative/Aki2-positive neurons subsequently die cell non-autonomously as neurodegeneration worsens. Transcriptomic analysis reveals a large number of genes dysregulated in the absence of neuronal Aki2, including a substantial subset that are downstream of the tumor suppressor p53. A role for p53 downstream of Aki2 is supported by a significant upregulation of p53 protein in knockout neurons, a similar dysregulation of p53 target genes in Aki2-null embryonic telencephalon transcriptomes, and a partial rescue of postnatal neurodegeneration when Aki2 mutants are heterozygous for p53. Together, these data identify a new role for Aki2 in postmitotic neuron maintenance, implicate p53 as a new functional partner underlying the pleiotropic roles played by the Akirin family of proteins, and suggest that Aki2 and its downstream partners may be relevant targets in a variety of neurodegenerative disorders.

## Results

### Neurons of the postnatal cerebral cortex express Akirin2

Multiple gene expression databases suggest that postmitotic neurons express *Aki2* transcript (Zhang et al., 2014; Mancarci et al., 2017; Sugino et al., 2019). To confirm postnatal protein and RNA expression, we performed immunofluorescence using an anti-Aki2 antibody and *in situ* hybridization with an antisense *Aki2* probe. Immunofluorescence (Figure 1A) and *in situ* hybridization (Figure S1A) of the adult mouse cortex revealed that cells in all cortical layers expressed Aki2 protein and transcript. Our previous work indicated that Aki2 transcripts were detectable in the mouse cerebral cortex from embryogenesis to early postnatal ages (Bosch et al., 2016), but dynamics of Aki2 protein expression throughout cortical development remained unknown. To assess this, we performed western blot analysis of cortical lysates at multiple postnatal ages using an anti-Aki2 antibody previously validated on knockout tissues (Bosch et al., 2018; Bosch et al., 2019). This revealed Aki2 expression peaked at P0 and tapered off during postnatal development. Cortical expression stabilized after three weeks of age and remained clearly detectable into adulthood (Figure 1B). Interestingly, mRNA levels assessed by qPCR at a sampling of ages remained consistent throughout development in contrast to Aki2 protein levels (Figure 1C). This is consistent with our prior work utilizing muscle cells as well as with a developmental mouse cortex transcriptomic dataset (Fertuzinhos et al., 2014; Bosch et al., 2019) and suggests that Aki2 protein is post-transcriptionally regulated during brain maturation. Furthermore, close inspection of western blots at a variety of exposures revealed 3 distinct bands of ∼22kDa, 25kDa, and 27kDa, as well as two higher molecular weight bands of ∼33kDa and ∼35kDa. Knockout and knockdown validation of the primary antibody used here (Bosch et al., 2018; Bosch et al., 2019), confirmed that all 5 of these bands are specific to Aki2. This suggests that the cortex expresses several post-translationally modified forms and/or unannotated Aki2 splice variants. The former appears more likely, given the mismatch between transcript and protein levels across ages and the lack of any splice variants upon examination of transcriptomic data and the *Aki2* genomic locus, as well as annotation in mouse genome browsers (data not shown).

**Figure 1:**
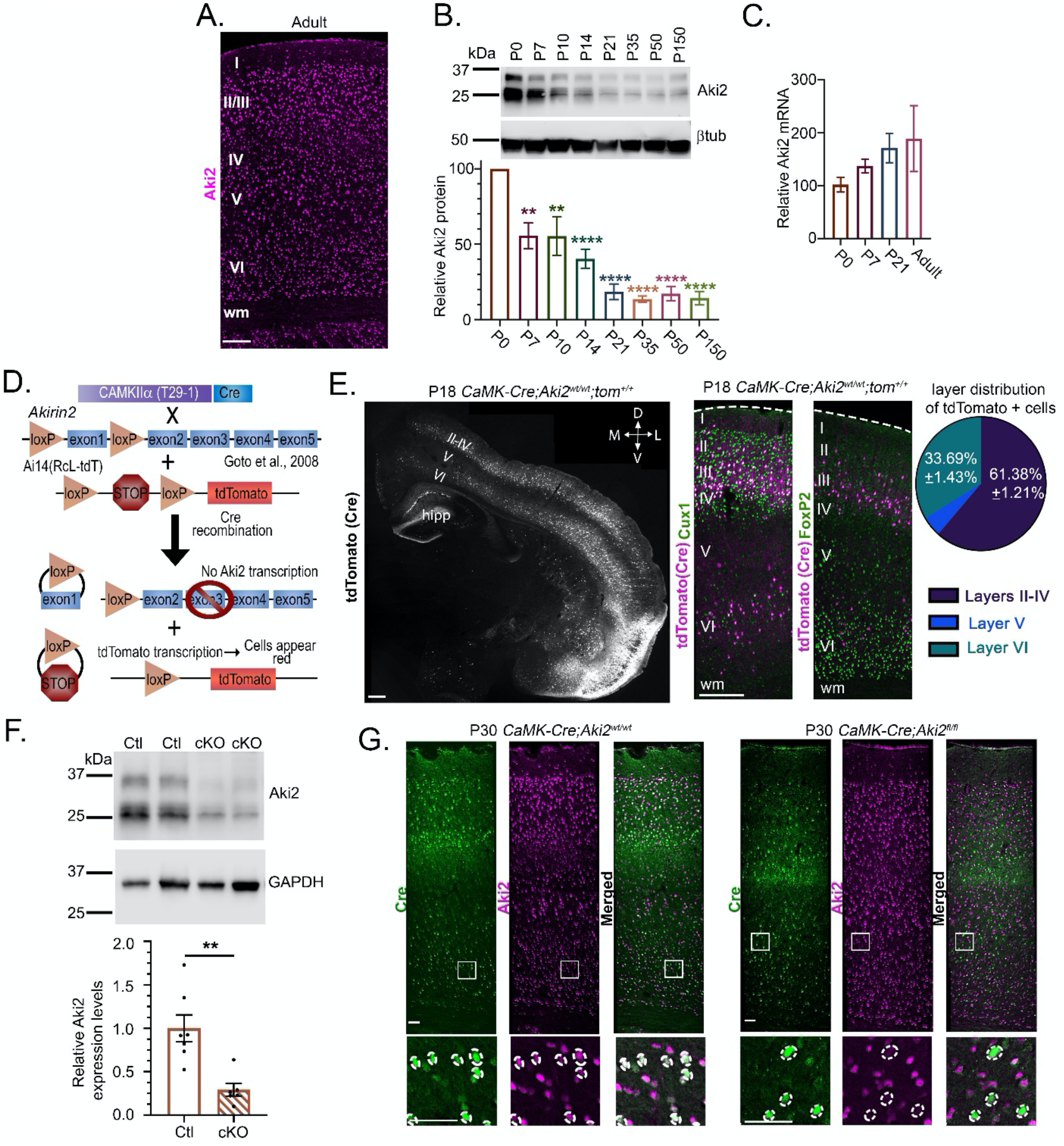
Generation of a restricted knockout mouse model to assess the role of Akirin2 during cortical maturation. A) Immunofluorescence of adult mouse cerebral cortex demonstrates widespread expression of Aki2 protein across all cortical layers (I-VI). B) Cortical lysates collected at the indicated ages were assessed via western blot and probed with an anti-Aki2 antibody. Multiple specific bands (see F) are observed around the expected molecular weight of 22 kDa, as well as two ∼33 kDa bands. Aki2 protein expression peaks around birth and slowly declines to a stable adult level of ∼20% relative to P0 (quantification of 3 samples per age, normalized to β−tubulin (βtub) signal within the same lane and graphed as percent of P0 expression level). C) Levels of Aki2 transcript do not change significantly across postnatal maturation of the cortex, indicating that Aki2 protein levels are post-transcriptionally regulated; qPCR of 3 samples per age, using GAPDH as a reference gene and plotted as percent of P0 levels. One-way ANOVA with Dunnet’s multiple comparisons test comparing every age to P0 was used for B and C. D) Schematic of mouse alleles and breeding strategy used. Mice expressing Cre under the CaMKIIα promoter (line T29-1; Tsien et al., 1996) were bred with mice in which exon 1 of *Aki2* is flanked by loxP sites, resulting in disruption of the *Aki2* locus in many excitatory neurons of the forebrain (Goto et al., 2008). The *Ai14-tdTomato* Cre reporter allele was introduced to label Cre-expressing cells. E) At P18, shortly after Cre expression onset, tdTomato labels many cells in cortical layers II-IV (Cux1-demarcated) and VI (FoxP2-demarcated), as well as CA1 and the molecular layer of the dentate gyrus of the hippocampus (hipp) in control mice. Quantification indicates that ∼2/3 of Cre-positive cells are located in layers II-IV and ∼1/3 are FoxP2-negative layer VI neurons; few, if any, Cre-positive cells are observed in layer V. F, G) Confirmation of Aki2 loss in *CaMK-Cre;Aki2*^*fl/fl*^ (cKO) mutant mice. F) Immunoblots of control and cKO cortical lysates with anti-Aki2 antibody indicate significant reduction of protein levels (and confirm specificity of all band sizes observed), with some protein expression remaining from Cre (-) cells. Values from 5-6 samples per genotype were normalized to GAPDH signal and compared using an unpaired t-test. G) Aki2 loss is confirmed at the cellular level using anti-Cre and anti-Aki2 antibody staining on cortical sections: Aki2 signal loss is observed in Cre-expressing cells in the mutant but not control cortex. D, dorsal; V, ventral; M, medial; L, lateral; wm, white matter; Scale bar: 300 µm in B, 100 µm in all other panels. Data shown as mean ± SEM, **p<0.01, ****p<0.0001. See also Figure S1.

### Selective *Akirin2* knockout in maturing cortical neurons

Consistent with its high levels of expression at embryonic ages, we previously found that Aki2 is essential for the normal proliferation, differentiation, and survival of progenitor cells throughout cortical development (Bosch et al., 2016). Given its maintained, albeit lower, expression in the postmitotic cortex, we hypothesized that Aki2 plays a distinct functional role in *postmitotic* neurons. To explore this role, we excised *Aki2* from excitatory forebrain neurons starting at ∼P18 by crossing mice harboring a conditional *Aki2* mutant allele (Goto et al., 2008) to the *CaMKIIα-Cre* line T29-1 (Tsien et al., 1996). We also included the *Ai14* reporter allele, which expresses the tdTomato red fluorescent protein from the ubiquitous ROSA locus when Cre excises a stop cassette flanked by loxP sites (Figure 1D). Using this reporter line, we found initial tdTomato expression spanning the somato-motor and somatosensory cortices in the rostral-caudal axis (Figure S1C). Although Cre was also active in hippocampal CA1, we limited our analysis to the cortex where close examination revealed that 61.38 ± 1.21% (mean ± SEM) of tdTomato-positive cells were found in layers II-IV and 33.69% ± 1.43% in layer VI, with very few in layer V (Figure 1E). There are two neuronal subpopulations in layer VI: FoxP2-negative corticocortical projection neurons and FoxP2-positive corticothalamic projection neurons (Kast et al., 2019). Close examination of layer VI revealed that most, if not all, of the FoxP2-positive corticothalamic neuron subpopulation lacked the Cre reporter; instead, Cre expression was limited to the subpopulation of FoxP2-negative corticocortical layer VI neurons (Figure 1E). Though it is possible that more cells activated *CaMK-Cre* expression as development proceeded, this was difficult to assess as tdTomato protein spreads to the neuropil, obfuscating individual cell bodies (Figure S1B). However, immunofluorescence using an anti-Cre antibody showed an expression pattern at P30 similar to that of tdTomato at P18 (Figure 1G), which is consistent with reported GFP expression driven by the CaMKIIα promoter (Wang et al., 2013). Altogether, these results indicate that in the *CaMK-Cre;Aki2*^*fl/fl*^ cortex, layers II-IV have the most Aki2-null neurons followed by FoxP2-negative neurons in layer VI, while layer V and FoxP2-positive neurons of layer VI are almost completely spared.

To confirm Aki2 loss, western blot of *CaMK-Cre;Aki2*^*fl/fl*^ and control cortical lysates was performed, revealing a ∼70% reduction of Aki2 protein (Figure 1F). As only a subpopulation of cortical cells expressed Cre and thus lost Aki2 expression in our model, the remaining signal from Cre-negative cells was expected. To ensure Aki2 loss is restricted to Cre-positive cells, P30 cortical tissue was immunostained with antibodies against Cre and Aki2. In *CaMK-Cre;Aki2*^*wt/wt*^ tissue, cells labeled with Cre still expressed Aki2, as expected. In *CaMK-Cre;Aki2*^*fl/fl*^ mutant tissues, Cre-positive cells lacked Aki2 staining whilst neighboring Cre-negative cells expressed Aki2 as normal (Figure 1G).

### Akirin2 mutants exhibit gradual neurodegeneration and cortical atrophy

*CaMK-Cre;Aki2*^*fl/fl*^ mutants were indistinguishable from littermate controls prior to, and for the first few weeks after, Aki2 loss. At ∼8 weeks of age, however, both male and female *CaMK-Cre;Aki2*^*fl/fl*^ animals stopped gaining weight. Males weighed significantly less than their littermate controls by 12 weeks of age, while females weighed significantly less by 18 weeks of age (Figure 2A). Mutants appeared to behave similarly to their control littermates until ∼P60, at which time they were often distinguishable by their hyperactivity and repetitive movements. We often found them incessantly running in circles or rearing up on their hindlimbs and scratching at the cage repetitively. At P150, the oldest age at which these mice were maintained, *CaMK-Cre;Aki2*^*fl/fl*^ mutants typically exhibited kyphosis (Figure 2B) and were largely inactive. Analysis of whole brains revealed a slow atrophy of the cortex while other Cre-negative structures (e.g., the cerebellum) remained unchanged (Figure 2C). To measure the degree of this atrophy, we measured cortical thickness (from lateral ventricle to pia) in coronal cryosections at each age. We found that in the absence of neuronal Aki2, the cortex was significantly thinner by P35, and continued to atrophy with age (Figure 2D). The thinning cortex was obvious in P50 Golgi-stained brains, where mutants consistently exhibited fewer labeled neurons (Figure 2E). Consistent with neuronal loss in the absence of Aki2, a sensitive silver stain for degenerating neurons (whose membranes become permeable) identified many impregnated neurons in mutants but not controls at P28 (Figure 2F), 10 days after Cre-onset and *Aki2* gene excision.

**Figure 2:**
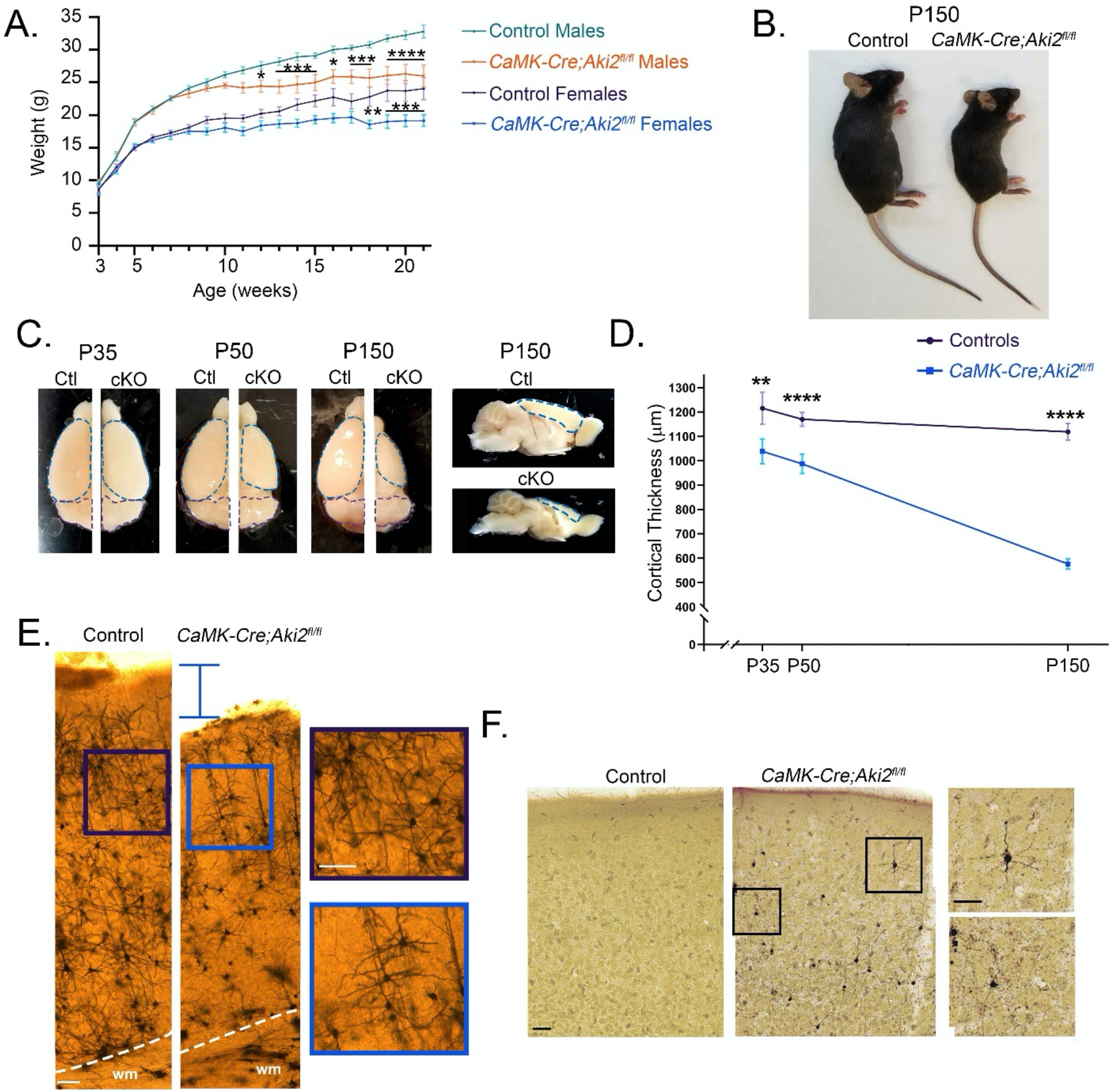
Severe, progressive cortical atrophy in the absence of neuronal Akirin2. A) Comparison of control mouse weights (n= 6-17 per time point) to weights of age and gender matched *CaMK-Cre;Aki2*^*fl/fl*^ mutants (n=4-13 per time point) indicates that mutant males and females stop gaining weight at ∼8 weeks of age. B) By P150 (∼21 weeks) *CaMK-Cre;Aki2*^*fl/fl*^ mutants are clearly much smaller and exhibit kyphosis. C) Whole brains of control (Ctl) and *CaMK-Cre;Aki2*^*fl/fl*^ (cKO) mice were collected at the indicated time points. The cortex is outlined in blue and the cerebellum in purple; the progressive atrophy of the cKO cortex (but not the Cre-negative cerebellum) is clear in comparison to controls. D) This atrophy was quantified by measuring cortical thickness from lateral ventricle to pia in cryosections of control and *CaMK-Cre;Aki2*^*fl/fl*^ mutant cortices (n=21-23 coronal sections from 4 mice per genotype). Data shown as mean ± SEM, *p<0.05, **p<0.01, ***p<0.001, ****p<0.0001, multiple t-tests with Holm-Sidak method corrections. E-F) Golgi (E) and NeuroSilver (F) staining on control and *CaMK-Cre;Aki2*^*fl/fl*^ cortices also reveal a thinner cortex (E; P50) with degenerating neurons (F; P28). Insets show magnified areas of the main image. Scale bar: 100 µm.

To determine the spatiotemporal extent of cell loss, we used NeuN immunofluorescence to label and estimate the quantity of all neurons in cortical sections (Figure 3A). The number of NeuN cells did not significantly differ between control and *CaMK-Cre;Aki2*^*fl/fl*^ cortices at P35. By P50, however, the *CaMK-Cre;Aki2*^*fl/fl*^ mutant cortex had significantly fewer neurons, and this loss continued through P90 and P150, by which point half of all neurons had been lost (Figure 3B). Cell loss differed by cortical layer in a manner consistent with Cre expression (Figure 1E). Neuronal loss was first apparent in layers II-IV, a population with many Aki2 null neurons, with significantly fewer cells remaining by P50 (Figure 3C, D), consistent with initial cell-autonomous death. In contrast, the number of neurons in subpopulations with sparse Cre expression (Layer V and FoxP2-positive layer VI) (Figure 1E) were roughly normal at P50 and did not significantly decrease until P90 (Figure 3D). This may indicate that later neuronal death can include cell non-autonomous effects of Aki2 loss in neighboring neurons.

**Figure 3:**
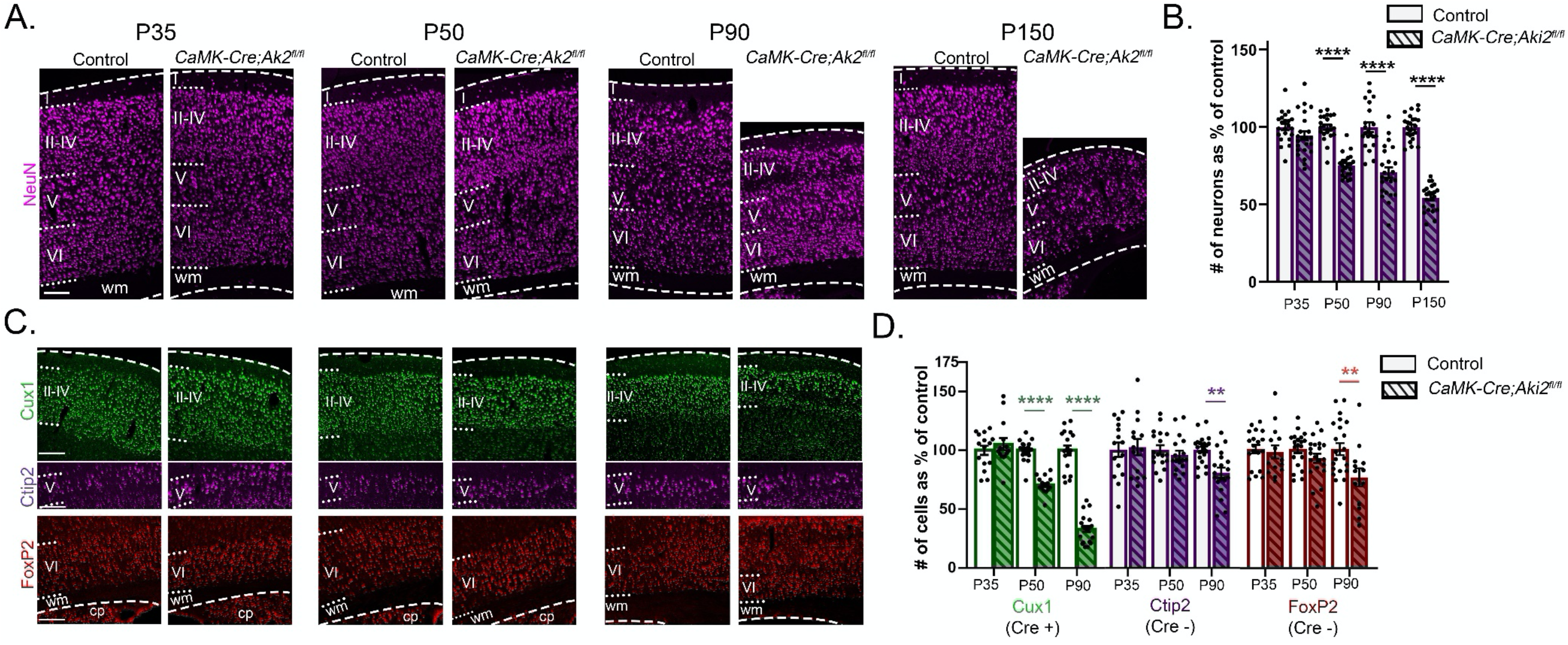
Massive cortical neuron loss in the absence of Akirin2. A,C) Cortical cryosections from control and *CaMK-Cre;Aki2*^*fl/f*l^ mice were immunostained for the pan-neuronal marker, NeuN (A) or the layer-specific neuronal markers (C) Cux1 (layers II-IV), Ctip2 (layer V), and FoxP2 (a subset of layer VI). B,D) The number of NeuN neurons in a layer-spanning strip of cortex (B) or the number of neurons belonging to each immunostained population (D) was quantified in 10-23 sections from 3-4 mice per genotype per age. Significant neuronal loss is observed by P50 and progresses with age (B). Cre-positive (and thus mutant) neuronal populations are lost first (D, Cux1), while Cre-negative neuronal populations (D, Ctip2, FoxP2) remain normal until P90. **p<0.01, ***p<0.001, ****p<0.0001 by 2-way ANOVA with Sidak’s multiple comparison tests comparing control to age-matched conditional knockout. Data shown as mean ± SEM. cp, caudoputamen; wm, white matter; dashed lines delineate cortical edges. Scale bar: 200 µm. See also Figure S2

Closer examination of layer VI allowed us to test the hypothesis that Cre-expressing (Aki2-null) neurons die cell-autonomously earlier than do Cre-negative (Aki2-positive) neurons. Immunofluorescence revealed that although 34% of Cre-expressing cells resided in layer VI, the vast majority of layer VI FoxP2-positive neurons were Cre-negative (Figure 1E). We thus used FoxP2-positive neurons as a proxy for Cre-negative neurons and FoxP2-negative neurons (quantified as the number of NeuN-positive/FoxP2-negative layer VI cells) as a proxy for Cre-positive neurons. Quantifications of these two populations revealed that although FoxP2-positive (Cre-negative) neuron numbers remained the same at P50 (Figure 3D), FoxP2-negative (Cre-positive) neuron numbers decreased by half by P50 (Figure S2). This is consistent with Aki2-null neurons dying cell-autonomously as an initial insult, between P35 and P50, followed by cell-non-autonomous death of neurons that retain Aki2 expression.

In response to neuronal injury or death, nearby astrocytes become activated and upregulate glial fibrillary acidic protein (GFAP) (Pekny et al., 2005). Additionally, microglia increase their phagocytic consumption of cellular debris resulting in an increase in microglial lysosomes (Brown et al., 2014). We asked whether activation of glial cells–all of which are Cre-negative and retain Aki2 expression (Zhang et al., 2014; Mancarci et al., 2017) in our model–occurred in the mutant cortex. We measured mean fluorescence intensity of GFAP antibody staining (Figure 4A) and the area immunostained with an antibody against CD68, a marker of lysosomes (Figure 4B). By P50, the GFAP mean fluorescence intensity was nearly 5-fold higher (Figure 4A), while CD68 immunolabeling increased by over 4-fold by P35 and 9-fold by P50, in the *CaMK-Cre;Aki2*^*fl/fl*^ mutant cortex (Figure 4B). Co-staining for CD68 and P2Y12, a microglial-specific receptor, confirmed that the increase in lysosomes was microglial-specific and demonstrated that microglia in the mutant cortex took on the characteristic amoeboid morphology of activated microglia (Figure 4B) (Stence et al., 2001). Therefore, concomitant with significant cell-autonomous neuronal loss, both astrocytes and microglia were activated in Aki2 mutants. Interestingly, we found that initially only layers II-IV and VI (where some neurons lose Aki2 expression) contained activated astrocytes and microglia while layer V (where Aki2 expression is unaltered) did not. Over time, however, activated astrocytes and microglia spread to layer V, concurrent with the secondary, likely cell-non-autonomous neuronal death observed at later ages.

**Figure 4:**
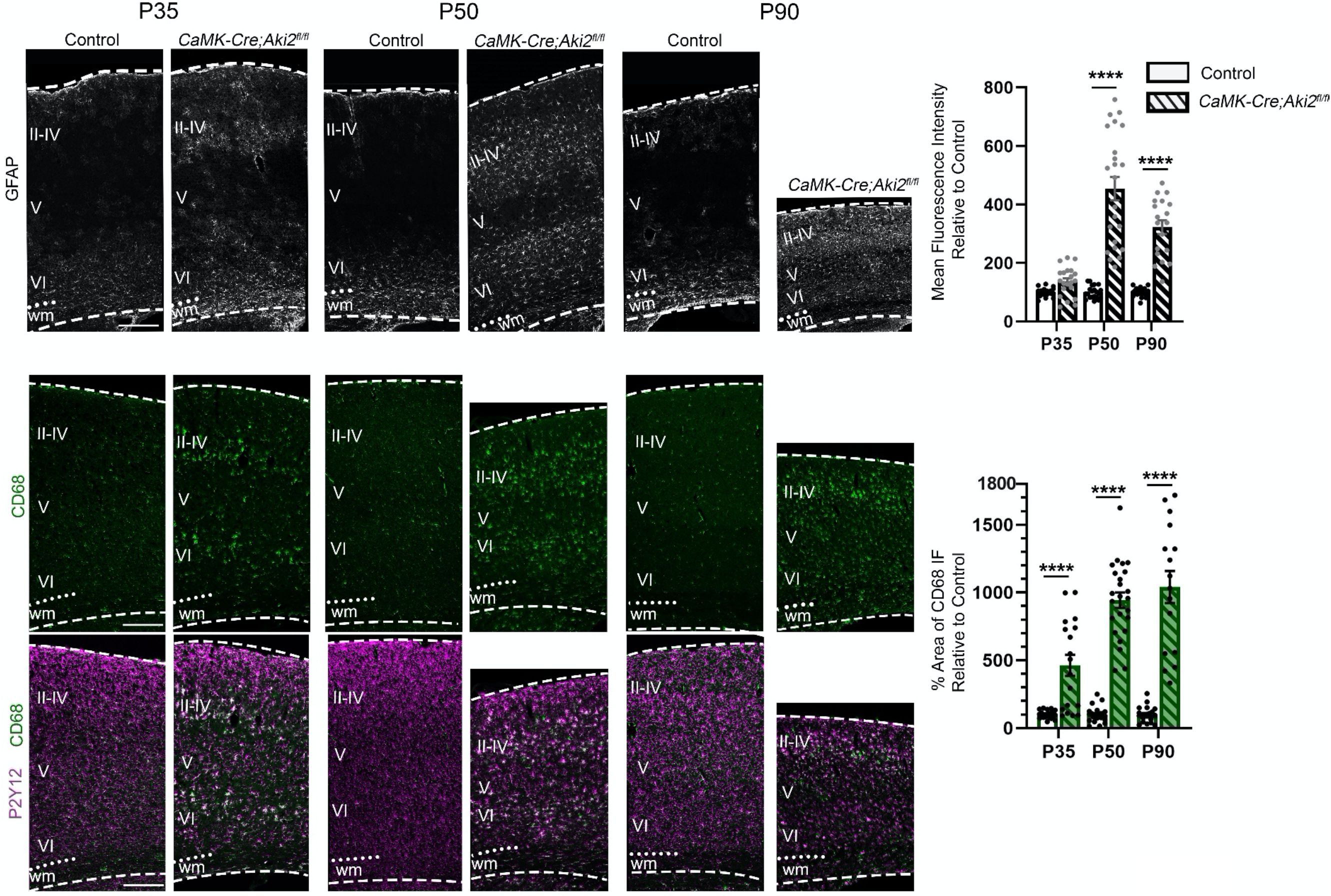
Glial activation accompanies cortical neuron loss in Akirin2 mutants. A) Control and *CaMK-Cre;Aki2*^*fl/fl*^ cryosections were immunostained with a marker for activated astrocytes (glial fibrillary acidic protein [GFAP]) or B) a marker for lysosomes (CD68) as a proxy for microglial phagic activity. B) Colocalization of CD68 staining with the microglial marker P2Y12 confirms activation of microglia. Fluorescence intensity measurements show remarkable increases in the activation of astrocytes (A) and microglia (B) across multiple time points, with the initial signal most prevalent in the layers containing significant numbers of dying Aki2-null neurons (II-IV and VI). ****p<0.0001 by 2-way ANOVA with Sidak’s multiple comparisons test comparing control to age-matched mutants. Data shown as mean ± SEM. n= 16-24 coronal sections from 3-4 mice per genotype per age. wm, white matter; dashed lines delineate cortical edges. Scale bar: 300 µm.

### Cortical neurons undergo necroptosis in the absence of Akirin2

Because previous studies found that Aki2 suppresses apoptosis in neural progenitor cells (Bosch et al., 2016), tumor cells (Krossa et al., 2015), and B lymphocytes (Tartey et al., 2015), we initially expected that the neuronal loss in *CaMK-Cre;Aki2*^*fl/fl*^ cortex would be apoptotic in nature. Surprisingly, neither immunofluorescence nor western blotting using an antibody for the apoptotic marker, cleaved caspase 3 (CC3), nor TUNEL staining for apoptotic nuclei, revealed any sign of excessive apoptosis in the mutant cortex at any of the ages examined (Figure S3A-C). In apoptosis, cells typically shrink, but we noticed that *CaMK-Cre;Aki2*^*fl/fl*^ cortical neurons, in contrast, appeared larger (Figure 5A). To confirm this, we measured NeuN labeling–which labels both the nucleus and the somatic cytoplasm (Lind et al., 2005) in a pattern that overlaps with Nissl stains (Figure S3D)–in *CaMK-Cre;Aki2*^*fl/fl*^ and control cortices at multiple ages. We found that *CaMK-Cre;Aki2*^*fl/fl*^ cortices contained larger neurons, and that this apparent cell swelling preceded cell death. In layers II-IV, where significant cell loss is first observed at P50, neurons were already larger at P35; in layers V and VI, where significant neuronal loss is not observed until P90, we found larger neuronal sizes beginning at P50 (Figure 5B).

**Figure 5:**
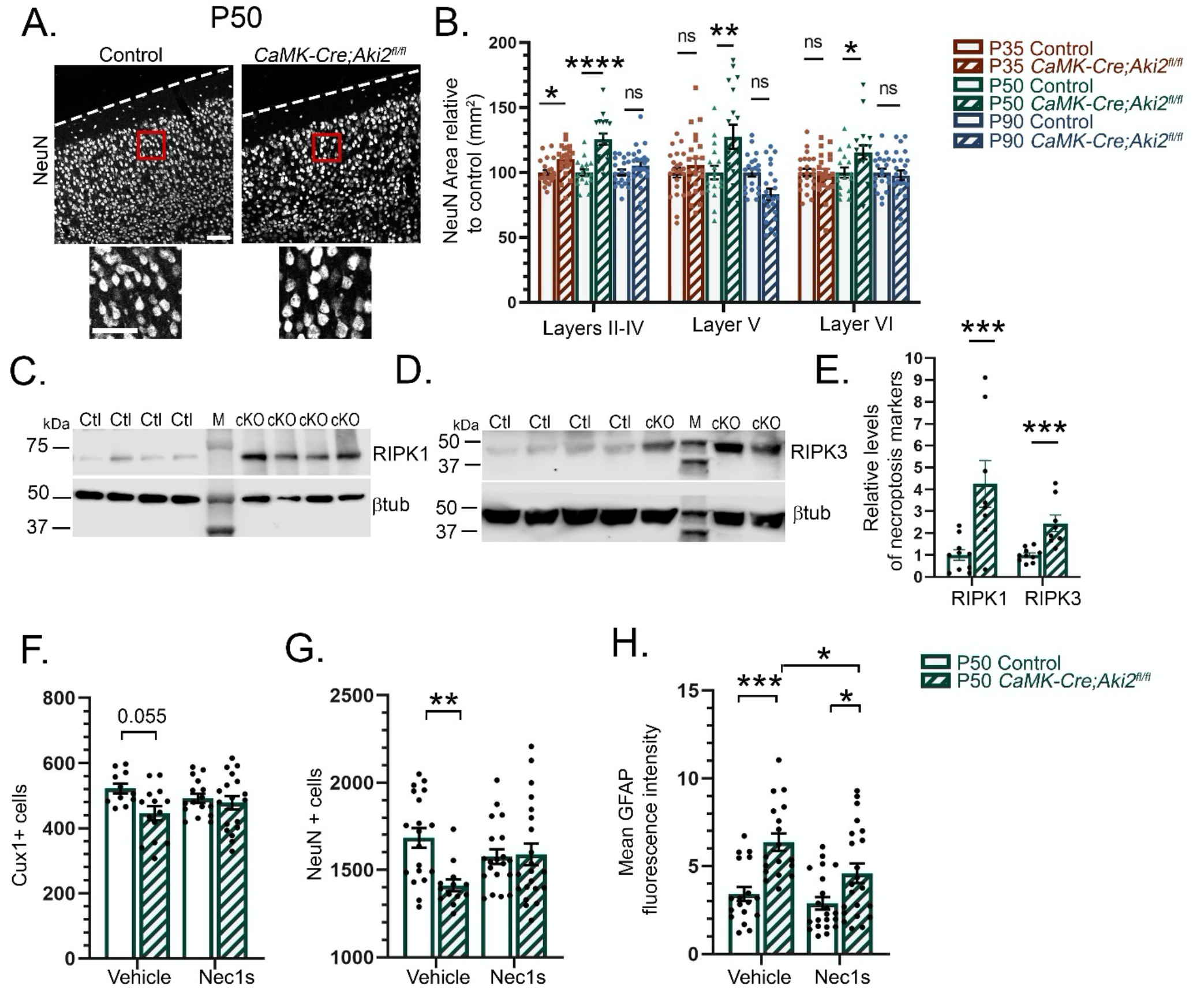
Cortical neurons undergo necroptosis in the absence of Akirin2. A) NeuN immunostaining reveals that *CaMK-Cre;Aki2*^*fl/fl*^ mutant neurons appear larger than control neurons. B) Neuronal size was estimated using NeuN area (nucleus+somatic cytoplasm) on 10 neurons per cryosection per layer (II-IV, V, VI) and the average size plotted as % control at each age indicated. Two-way ANOVA with Sidak’s multiple comparisons tests comparing control to age-matched conditional knockout. C-E) Cortical lysates of P50 control (n=10) and *CaMK-Cre;Aki2*^*fl/fl*^ (cKO) mice (n=8) were immunoprobed for necroptosis-regulating kinases RIPK1 (C) and RIPK3 (D); M= molecular weight marker. E) Levels of the indicated protein were normalized to β-tubulin within the same lane, graphed as percent of control, and statistical significance determined using the Holm-Sidak method for 2 unpaired t-tests. Both necroptosis markers are significantly increased in Aki2 mutant cortex. F-H) Mice were injected daily with vehicle (DMSO) or the necroptosis inhibitor Nec-1s for 30 days. The ability of Nec1s to rescue neuronal death was assessed using immunofluorescence for Cux1 (upper layer neurons; F) or NeuN (all neurons; G). Glial activation was also assessed by measuring GFAP fluorescence intensity (H). Statistical significance was determined using 2-way ANOVA with Tukey’s multiple comparison test comparing all means regardless of treatment and genotype. Daily Nec-1s treatment significantly rescued neuronal number and degree of glial activation in Aki2 mutant cortex. n= 11-21 coronal sections from 3-4 mice per condition. Data shown as mean ± SEM. *p<0.05; **p<0.01; ***p<0.001; ****p<0.0001. βtub, β−tubulin; Scale bar: 150 µm. See also Figure S3.

An increase in cell size can be indicative of either autophagy or necroptosis (Kroemer et al., 2009). To determine if Aki2-null cortical neurons were undergoing autophagy, we utilized western blot assays to examine the lipidation of the microtubule-associated protein 1 light chain 3 (LC3) from LC3-I to LC3-II, a key step in the formation and lengthening of autophagosomes (Kabeya et al., 2000). We found no increase in the faster-migrating LC3II band vs. the LC3I band in *CaMK-Cre;Aki2*^*fl/fl*^ cortical lysates compared with controls (Figure S3E), making autophagy an unlikely explanation for the neuronal death we have observed. In contrast, levels of necroptosis mediators, receptor-interacting protein kinases RIPK1 and RIPK3, were robustly and consistently increased in *CaMK-Cre;Aki2*^*fl/fl*^ cortical lysates (Figure 5C-E), suggesting that neurons undergo necroptosis in the absence of Aki2. To provide further support for this hypothesis, we injected a cohort of mice daily with the RIPK1-specific inhibitor, Nec-1s, or vehicle from P21 (shortly after Cre expression begins and the Aki2 allele is excised; Figure 1) until P50 (the age at which we first detect significant neuronal loss; Figure 3). We then assessed cell numbers and glial activation levels to determine if application of this necroptosis inhibitor ameliorated the phenotypes seen in *CaMK-Cre;Aki2*^*fl/fl*^ mutant cortex. We found that *CaMK-Cre;Aki2*^*fl/fl*^ mutants that received Nec-1s and controls had comparable numbers of Cux1-positive Layer II-IV neurons (Figure 5F) and total cortical neurons (Figure 5G), while mutants receiving vehicle exhibited the expected deficits. GFAP levels were significantly increased in both vehicle-treated and Nec-1s-treated *CaMK-Cre;Aki2*^*fl/fl*^ mutants compared to their controls, but less so in Nec-1s-treated *CaMK-Cre;Aki2*^*fl/fl*^ mutants than in mutants treated with vehicle only (Figure 5H). Together, we interpret these results as a partial rescue of the Aki2 mutant phenotype by Nec-1s, which bolsters the other evidence–lack of apoptosis, lack of evidence for autophagy, upregulation of RIPK1/3, and neuronal swelling–indicating that cortical neurons lacking Aki2 die over a period of several months by necroptosis.

### Akirin2 loss disrupts gene expression patterns in both the postnatal and embryonic cortex

Aki2 regulates gene expression patterns in many organisms in a cell type- and context-dependent manner (Komiya et al., 2008; Nowak et al., 2012; Akiyama et al., 2013; Bonnay et al., 2014; Komiya et al., 2014; Tartey et al., 2014; Bosch et al., 2020). We thus hypothesized that Aki2 regulates overlapping, but distinct, gene patterns throughout the generation and maturation of the cortical neuronal lineage. To test this, we performed transcriptomic analyses on cortical tissues using two different conditional Aki2 knockout models: 1) the *CaMK-Cre;Aki2*^*fl/fl*^ model described here, in which the Aki2 allele is disrupted in cortical neurons starting ∼P18; and 2) the *Emx1-Cre;Aki2*^*fl/fl*^ model described in Bosch et al. (2016), in which the Aki2 allele is disrupted in cortical ventricular zone neural progenitors starting at E9.5. We did this for two reasons: 1) to generate hypotheses about what biological processes are regulated by Aki2 in the developing and maturing brain, a topic that remains unexplored; and 2) to compare and contrast the embryonic vs. postnatal differentially expressed genes (DEGs) to derive common, as well as context-specific, Aki2-dependent genes.

We performed RNA sequencing (RNASeq) on cortical tissue from *CaMK-Cre;Aki2*^*fl/fl*^ mice and their littermate controls at P35, as this allowed substantial time for existing Aki2 protein turnover after Cre onset but was substantially prior to obvious neuronal loss (P50). Transcriptomic analysis using an FDR cut-off of 0.05 identified 295 genes significantly upregulated and 156 genes significantly downregulated when Aki2 is lost from many cortical neurons (Figure 6A). Gene set enrichment analysis (GSEA) identified vesicle targeting as the top overrepresented gene ontology (GO) biological process as well as several neuronal-specific GO biological processes (regulation of postsynaptic membrane potential, regulation of membrane potential, action potential propagation), and several pathways involved in regulating the mitotic cell cycle, which would be aberrant in postmitotic neurons (Figure 6B,D).

**Figure 6:**
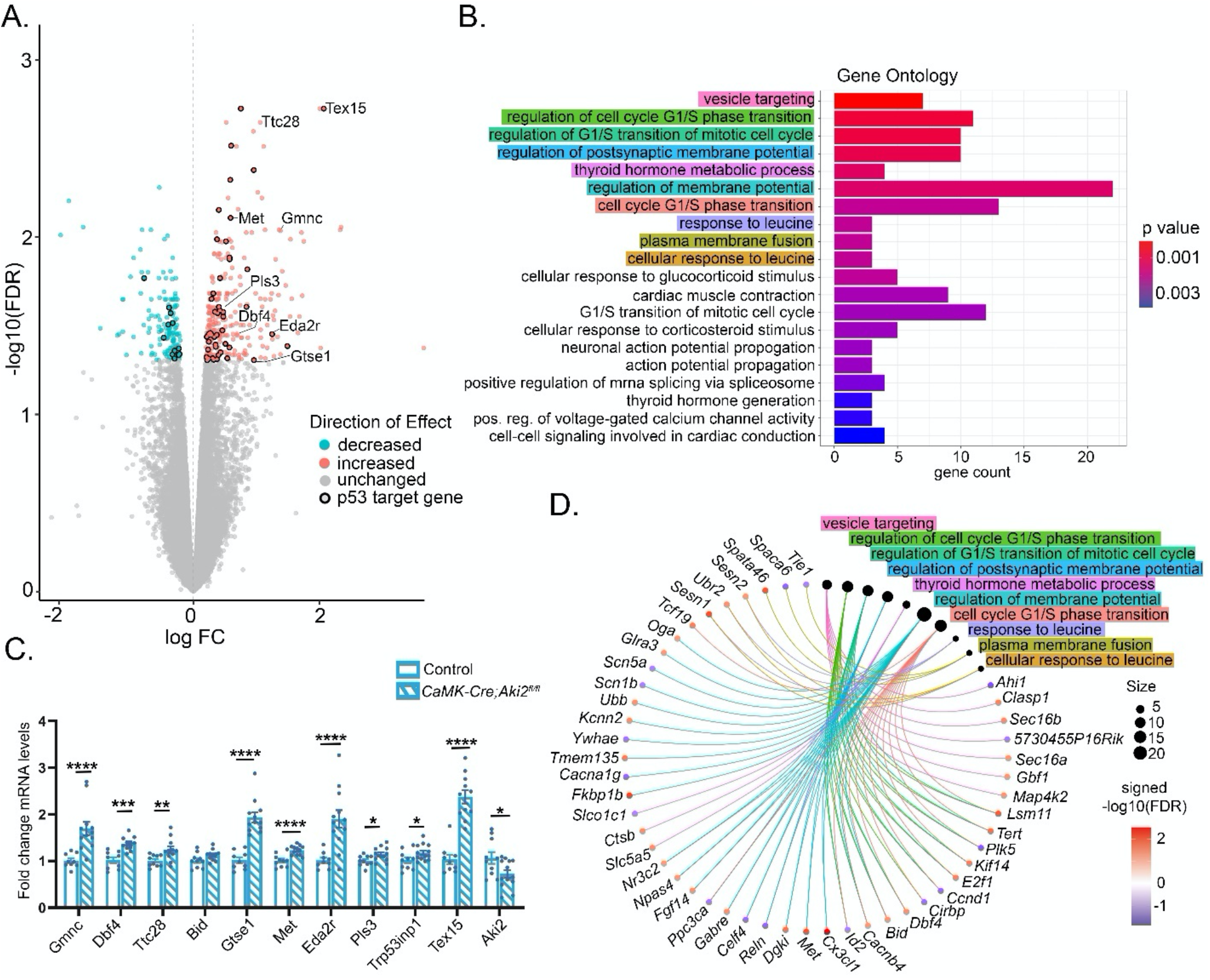
Akirin2 loss disrupts gene expression patterns in the postnatal cortex. RNA was extracted from cortical tissues of P35 control (n=4) and *CaMK-Cre;Aki2*^*fl/f*l^ mice (n=3) and analyzed by RNASeq and a bioinformatic pipeline as described in Materials and Methods. A) Volcano plot showing genes whose expression is significantly upregulated (red) or downregulated (blue) in Aki2 mutant cortex with an FDR cutoff of 0.05. Genes in the p53 pathway are highlighted with black circles. B) Gene ontology (GO) analysis bar plot showing biological processes associated with the differentially expressed genes (DEGS) identified in (A). C) qPCR Confirmation of upregulated cell-cycle and p53 target genes in Aki2 mutant biological and technical replicates (n=10-13) compared to control (n=8-10). Note confirmation of significant (but not complete) Aki2 downregulation as expected. D) Gene-concept network of the GO biological processes showing the relationship between the GO terms highlighted in B and the individual genes comprising each group. Size of the black circle for each GO term indicates the number of included genes, while color of individual gene circles corresponds to the signed - log10(FDR) as indicated in the legend. Data shown as mean ± SEM. FC, fold change; FDR, false discovery rate. See also Figure S4 and Tables S3-7

To gather comparative data on embryonic cortical neural progenitors, we also performed RNASeq on dorsal telencephalic tissue from E10.5 *Emx1-Cre;Aki2*^*fl/fl*^ mice and their littermate controls. As above, we carefully chose this time point as one where we can detect Aki2 loss, but at which significant apoptotic cell death has not yet occurred (Bosch et al., 2016). Using a 0.05 FDR cut-off, 49 genes were significantly upregulated while 16 genes were significantly downregulated (Figure S4A). Interestingly, GSEA revealed that these DEGs are involved in the p53-mediated apoptotic signaling pathway and in negatively regulating the transforming growth factor beta (TGF-β) receptor signaling pathway, in which the Aki2 *C. elegans* orthologue, Akirin, participates (Bowman et al., 2020) (Figure S4B,D). Using this conditional knockout mouse model, we previously found that Aki2 is essential for dorsal forebrain development because it controls cell cycle progression and suppresses apoptosis in cortical neural progenitor cells (Bosch et al., 2016). While GSEA of DEGs with an FDR cutoff of 0.05 initially identified the GO term hindbrain development, the individual genes involved (Figure S4D), *tbr1* (McKenna et al., 2011; Fazel Darbandi et al., 2018), *lmx1b* (Donovan et al., 2019), *wnt7a* (Qu et al., 2013), (Fernando et al., 2014), *fzd4* (Bian et al., 2015), *zfp365* (Hattori et al., 2007; Okamoto et al., 2015) are involved in neurogenesis and/or neuron development in many brain regions, not exclusive to the hindbrain. Furthermore, relaxing the FDR cut-off to 0.1 results in a gene list enriched in DEGs associated with GO terms such as forebrain development, cell fate specification, and cell fate commitment (Figure S4C), further supporting Aki2’s role in these processes as found earlier (Bosch et al., 2016; Liu et al., 2017).

To identify potential Aki2-dependent genes across multiple cell types and stages, we compared DEGs across both transcriptomes. We identified five genes (*Trp53inp1, Ccng1, Eda2r, Susd6, Pls3*) that were upregulated and one gene (*Espn*) that was downregulated in both datasets (Table S3). Comparing the KEGG pathways enriched in both transcriptomes revealed that they both converge on the p53 signaling pathway (Figure S4F). In fact, five of the six DEGs found in both transcriptomes are p53 target genes. The p53 protein, encoded by the *Trp53* gene, is a transcription factor that responds to cellular stress to alter transcription of genes regulating a wide array of biological processes including necroptosis (Wang et al., 2016), apoptosis, cell cycle arrest, and proliferation(Miller et al., 2000; Murray-Zmijewski et al., 2008). Interestingly, using Ingenuity Pathway Analysis (IPA, Qiagen) (Krämer et al., 2013) to predict upstream transcriptional regulators of the two transcriptomes, we found that p53 was the most highly significant predicted regulator for both (Tables S4 & S5). p53 regulates 56 DEGs in the neuronal transcriptome (Table S6 and black outlined data points in Figure 6A), and 23 DEGs in the neural progenitor transcriptome (Table S7 and black outlined dots in Figure S4A).

Furthermore, several p53-linked biological processes (proliferation, differentiation, and apoptosis in *Emx1-Cre;Aki2*^*fl/fl*^ mice and neurodegeneration and necroptosis in *CaMK-Cre;Aki2*^*fl/fl*^ mice) are dysregulated in our mutant mouse models (as shown above and in Bosch et al., 2016). Additionally, p53-linked GO terms in each transcriptome (apoptotic signaling, response to UV, and fibroblast proliferation in *Emx1-Cre;Aki2*^*fl/fl*^ transcriptome and cell cycle transition in *CaMK-Cre;Aki2*^*fl/fl*^ transcriptome) further support aberrant p53 signaling in both Aki2 conditional mutants. We subsequently performed qPCR validation experiments choosing, based on preliminary DAVID (Huang da et al., 2009a; Huang da et al., 2009b) and IPA analysis, several p53 target genes and 3 Aki2-dependent genes common to both transcriptomes. Because Liu et al., 2017 report Aki2 binding to Geminin in *Xenopus*, we also validated the purported upregulation of a geminin family member, *Gmnc* in the *CaMK-Cre;Aki2*^*fl/fl*^ transcriptome. This independent analysis confirmed significant upregulation of 9/10 genes analyzed in *CaMK-Cre;Aki2*^*fl/fl*^ biological and technical replicates including *Gmnc* (p=0.0001), *Dbf4* (p=0.0016), *Ttc28* (p=0.0100), *Gtse1* (p<0.0001), *Met* (p=0.0003), *Eda2r* (p=0.0009), *Pls3* (p=0.0243), *Trp53inp1* (p=0.0409), and *Tex15* (p<0.0001) (Figure 6C). In *Emx1-Cre;Aki2*^*fl/fl*^ biological replicates, we confirmed that *Ano3* (p=0.0056), *Cdkn1a* (p=0.0030), and *Trp53inp1* (p=0.01) were significantly upregulated, that *Aki2* (p=0.0096) was significantly reduced as expected, and that *Eda2*r (p=0.0622) was upregulated though did not reach significance (Figure S4E).

### Akirin2 inhibits P53-mediated cell death

The *CaMK-Cre;Aki2*^*fl/fl*^ transcriptome and qPCR confirmation experiments (Figure 7A) indicated that *Trp53* transcript levels did not change significantly in the mutant, yet many p53 target genes were significantly upregulated. This is consistent with the fact that p53 is regulated primarily through post-translational modifications (PTMs), and not by changes in transcript expression (Oren 1999). For this reason, we measured p53 protein levels in cortical lysates and discovered a significant and unequivocal increase in *CaMK-Cre;Aki2*^*fl/fl*^ mutants (Figure 7B). Furthermore, immunostaining of *CaMK-Cre;Aki2*^*fl/fl*^ cortical sections revealed that only Aki2-null neurons exhibited a cell-autonomous massive upregulation of p53 protein levels (Figure 7C). Altogether, this suggests that Aki2 suppresses p53 activation in the adult cortex. Because p53 is a major regulator of cell viability (Morrison et al., 2000; Ranjan et al., 2016), we tested the hypothesis that p53 mediates the neurodegeneration we have documented in the *CaMK-Cre;Aki2*^*fl/fl*^ cortex. We crossed *CaMK-Cre;Aki2*^*fl/fl*^ mice to p53 constitutive knockout mice (Jacks et al., 1994) to generate p53 heterozygous Aki2 mutants (*CaMK-Cre;Aki2*^*fl/fl*^ ;*Trp53+/-*). If high p53 levels cause cell death, we would expect that reducing those levels (by removing one allele) would rescue neuronal death and subsequent glial activation in *CaMK-Cre;Aki2*^*fl/fl*^ mice; such haploinsufficiency has been observed previously for p53-mediated cell death (Sakhi et al., 1996; Hirata et al., 1997; Aloyz et al., 1998; Bae et al., 2005; Ghosh et al., 2018). Indeed, we found a significant rescue: *CaMK-Cre;Aki2*^*fl/fl*^ ;*Trp53+/-* mice had normal neuronal numbers (Figure 7D) and reduced glial activation compared to *CaMK-Cre+;Aki2*^*fl/fl*^ mice (Figure 7E-F). Together, these data indicate that the absence of Aki2 in postmitotic neurons leads to post-transcriptional upregulation of p53 and consequent neurodegeneration that requires p53-mediated pathways.

**Figure 7:**
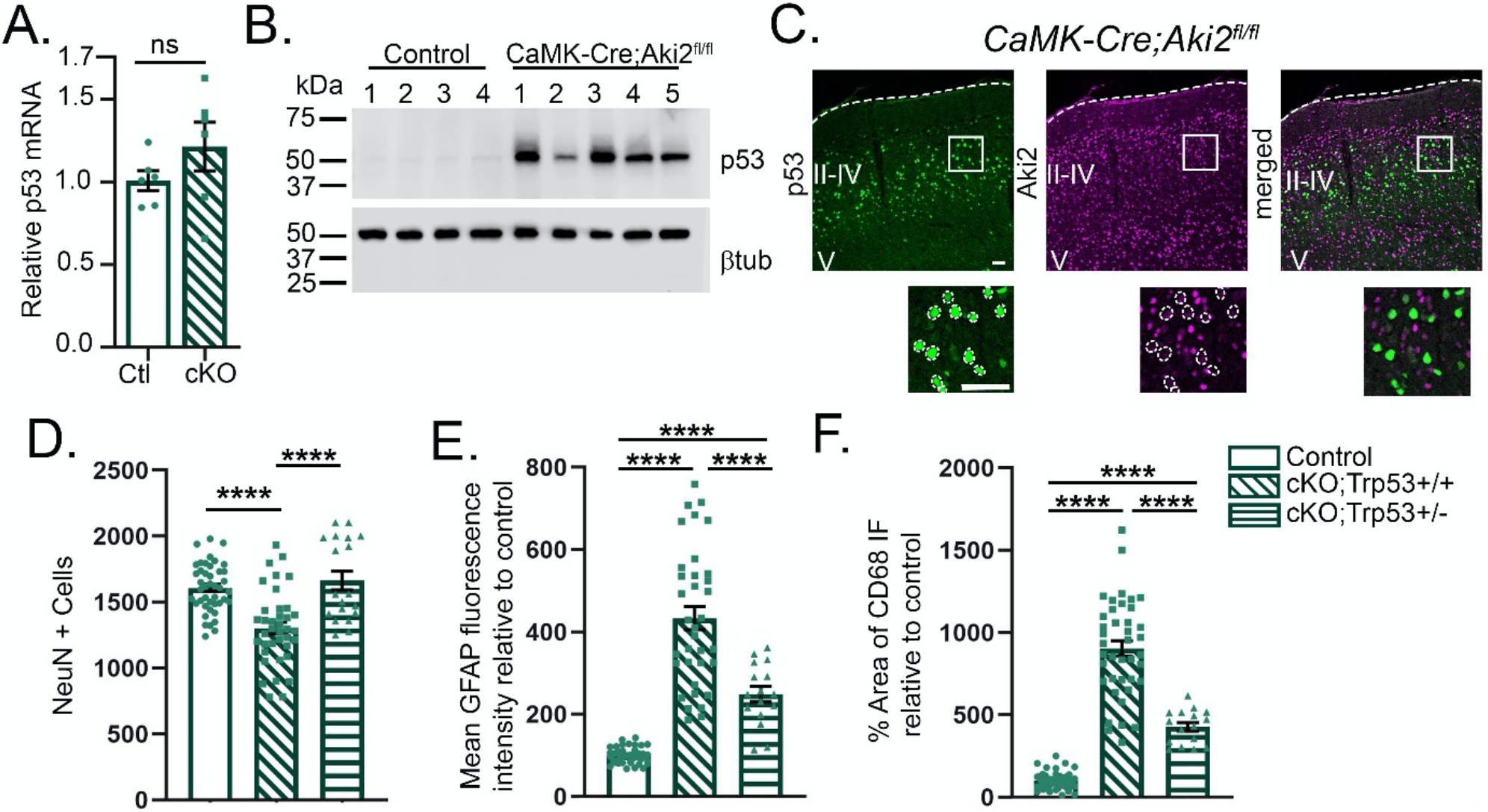
Neurodegeneration in Akirin2 mutant cortex is mediated by p53. p53 protein, but not transcript, is upregulated in Aki2 null neurons. A) qPCR using p53 primers identifies no significant upregulation of p53 transcription in the Aki2 mutant cortex (n=12) compared to control (n=10). Significance determined using unpaired t-test. B,C) In contrast, western blots (B) and immunostaining of cortical cryosections (C) using an anti-p53 antibody reveal a massive increase in protein levels restricted to Aki2 null neurons (dashed circles, layers II-IV). D-F) p53 heterozygosity largely rescues signs of neurodegeneration in Aki2 mutant cortex. Cryosections were stained with antibodies against NeuN, GFAP, and CD68 and quantified as in previous figures. Data for control and cKO;Trp53+/+ were replotted from Figure 3 with the addition of 18-21 sections from 2-4 newly analyzed animals that were littermates to the cKO;Trp53+/- (n=19 sections from 3 animals). Reduction in p53 levels completely rescues neuronal numbers (D) and partially rescues aberrant glial activation (E,F). βtub, β−tubulin. Data shown as mean ± SEM. ****p<0.0001. Statistical significance was determined using one-way ANOVA with Tukey’s multiple comparison test comparing genotypes. Scale bar: 100 µm.

## Discussion

The small, nuclear protein, Aki2, is broadly expressed in an array of tissues where its most consistently reported functions include the control of cell proliferation and differentiation (Bosch et al., 2020). The expression of Aki2 in non-proliferating cells such as maturing postnatal neurons suggested additional roles. Here, we have used the CaMKIIα-Cre driver to selectively remove Aki2 from many postmitotic neurons of the cerebral cortex at ∼P18. Following Aki2 excision, neurons (and animals) undergo a progressive decline in health, beginning with positive silver staining and cortical thinning without significant cell loss, suggestive of initial neurite degradation and neuropil depletion. By P50, ∼30 days after Aki2 loss, mutants exhibit significant neuron loss and cortical atrophy that worsens with age. Cell death and glial activation moves from the cortical layers with the greatest number of Aki2 null cells and progresses to primarily Cre-negative layers providing evidence that primary cell death occurs in Aki2-null neurons cell-autonomously followed by secondary cell-non-autonomous death in neurons that retain Aki2. Interestingly, unlike the apoptosis reported in embryonic Aki2-null cortical progenitors (Bosch et al., 2016) and muscle progenitors (Bosch et al., 2019) we present extensive convergent evidence that Aki2-null postmitotic neurons die via necroptosis in a p53-dependent manner. Our transcriptomic analyses also support this conclusion, as both mutant neural progenitor and postmitotic neuron transcriptomes were enriched for upregulated p53 target genes. Our data implicate p53 pathway proteins as novel mediators of Aki2 function and suggest that mutation and/or dysregulation of Aki2 and its partners could contribute to a variety of neurodegenerative disorders.

### Neurodegeneration in *CaMK-Cre;Aki2*^*fl/fl*^ mice: possible mechanisms

GSEA of the *CaMK-Cre;Aki2*^*fl/fl*^ (postmitotic cortical neuron-knockout) transcriptome reveals dysregulation of genes controlling the cell cycle. This is striking, because neurons are terminally differentiated cells that must remain in the mitotically quiescent G_0_ phase of the cell cycle. Cyclin-dependent kinases (Cdks) together with cyclins regulate a cell’s progression through the stages of the cell cycle (Boward et al., 2016). Numerous studies reveal aberrant expression of multiple cyclins, Cdks and Cdk inhibitors in several neurodegenerative diseases. Together with induction of signal transduction pathways and cytoskeletal changes reminiscent of developing neurons, this suggests that neurons in neurodegenerative tissues aberrantly re-enter the cell cycle (Zhu et al., 2007). Because neurons are ill-equipped for cell division, this is believed to induce neuronal death (Park et al., 2007) as there is a lack of evidence for successful division in postmitotic neurons (Zhu et al., 2007). The *CaMK-Cre;Aki2*^*fl/fl*^ mutant transcriptome is enriched in genes that regulate the G1/S mitotic transition, including several cyclins and cyclin dependent kinases.

Of particular interest to this study, the retinoblastoma tumor suppressor protein (pRb) suppresses the G1 to S phase transition in a signaling cascade involving D-type cyclins, Cdk4/6, and the E2F protein family. In quiescent neurons, hypophosphorylated pRb represses members of the E2F protein family to block cell cycle progression (Lees et al., 1993). In response to a mitogenic stimulant, D-type cyclins are synthesized and bind to Cdk4 or Cdk6. The cyclin D-Cdk4/6 complex phosphorylates and represses pRb, leading to E2F activation (Giacinti et al., 2006; Narasimha et al., 2014). Increased E2F-1 can result in G1/S cycle transition (Qin et al., 1994) or cell death (Hsieh et al., 1997; MacManus et al., 2003; Polager et al., 2008). Interestingly, *E2f1* and *Ccnd1* (encoding cyclin D1) are both increased in *CaMK-Cre;Aki2*^*fl/fl*^ cortices. It is thus possible that Aki2 represses cyclinD1; in Aki2 mutants, cyclinD1 may transactivate E2F-1 and trigger cell cycle re-entry and death. Together, this suggests that Aki2 maintains postmitotic neuronal health by suppressing expression of aberrant mitotic genes.

This would be in contrast to Aki2 functions in proliferating cells, where Aki2 *activates* expression of cell cycle genes. Previous studies suggest that Aki2 induces cyclin D1 and cyclin D2 expression and is critical for recruiting the chromatin remodeling complex containing Brg1 to the promoter of cyclin D2 in proliferating B cells (Tartey et al., 2015). This is further evidence supporting the hypothesis that Aki2 has cell-type- and context-dependent functions that are specific to each developmental stage.

### P53 as a mediator of Akirin2 mutant phenotypes

Because inactivation of the tumor suppressor protein, p53, is found in almost half of all human cancers (Kandoth et al., 2013; Lawrence et al., 2014), there has been extensive focus on its ability to drive cell cycle arrest and induce apoptosis. The p53 protein, however, is also involved in many other biological processes. Studies suggest that p53 hyperactivation can explain the neurodegeneration and necroptosis we show here in *CaMK-Cre;Aki2*^*fl/fl*^ mutants. Reminiscent of the massively increased p53 protein levels in Aki2-null cortical neurons, increased p53 levels are also found in relevant areas of the central nervous system in patients with neurodegenerative diseases, as well as in mouse models of such disorders (Miller et al., 2000). Additionally, neurons upregulate p53 in response to insults such as excitotoxicity (Morrison et al., 1996; Xiang et al., 1996), ischemia (Banasiak et al., 1998; Watanabe et al., 1999), traumatic brain injury (Plesnila et al., 2007), and neurotoxicity (Hirata et al., 1997; Morrison et al., 2000). In mouse models, ablation or reduction of p53 prevented cell death and/or rescued behavioral deficits (Herzog et al., 1998; Morrison et al., 2000; Bae et al., 2005; Mfossa et al., 2020). The upregulation of several p53 pathway genes in Aki2-null cortex and our observation that heterozygous p53 mutation rescued neuronal death in *CaMK-Cre; Aki2*^*fl/fl*^ mice suggest that dysregulation of an Aki2-p53 axis is worth considering as a potential contributor to the progression of neurodegenerative disorders.

Although p53 activity is largely associated with apoptosis, previous work has identified p53 involvement in autophagy (Maiuri et al., 2010) and regulated forms of necrosis including both necroptosis and ferroptosis (Tu et al., 2009; Vaseva et al., 2012; Jiang et al., 2015; Ranjan et al., 2016; Wang et al., 2016). p53 contributes to RIPK1/RIPK3-mediated necroptosis by upregulating levels of a long non-coding RNA (lncRNA) called necrosis-related factor (NRF). NRF inhibits miR-873, which in turn inhibits RIPK1 and RIPK3, the essential necroptosis initiators. Thus, upregulation of p53 leads to upregulation of NRF, increasing inhibition of miR-873 and disinhibiting RIPK1 and RIPK3 resulting in necroptosis (Ranjan et al., 2016; Wang et al., 2016). Admittedly, neither NRF nor miR-873 were significant DEGs in the *CaMK-Cre;Aki2*^*fl/fl*^ transcriptome. This may be because RNASeq was performed at P35, before obvious neuronal loss, or that Cre activity is not ubiquitous in this model and thus many neurons retain normal Aki2 expression. In any case, our results are altogether consistent with p53 activation leading to necroptosis in *CaMK-Cre;Aki2*^*fl/fl*^ mutant cortex.

Significant enrichment in p53-target genes in the *Emx1-Cre;Aki2*^*fl/fl*^ transcriptome further support the hypothesis that p53 mediates Aki2 function as p53 functions can explain the embryonic phenotypes we previously reported (Bosch et al., 2016). In the developing nervous system, p53 regulates neural stem cell proliferation and neural progenitor differentiation (Ferreira et al., 1996; Armesilla-Diaz et al., 2009; Forsberg et al., 2013; Quintens et al., 2015; Mfossa et al., 2020), as does Aki2 (Bosch et al., 2016; Liu et al., 2017). Interestingly, a recent preprint by Mfossa et al. (2020) revealed that p53 hyperactivation via gamma irradiation at E11 disrupts the cell cycle, triggers apoptosis, and leads to premature neurogenesis and ectopic neurons in the subventricular zone within 24 hours of irradiation. This results in microcephaly in newborn pups and is partially rescued by *Emx1-Cre*-mediated *Trp53* ablation in neural progenitor cells (Mfossa et al., 2020). Furthermore, adherens junctions lining the ventricular surface were disrupted and several adherens junction proteins were downregulated after p53 activation (Mfossa et al., 2020). These aberrations closely resemble those found in the *Emx1-Cre;Aki2*^*fl/fl*^ telencephalon (Bosch et al., 2016). Altogether, this implies that the Aki2-null cortical progenitor phenotypes (Bosch et al., 2016) result, at least in part, from hyperactivation of p53. *Emx1-Cre;Aki2*^*fl/fl*^ mice displayed more severe microcephaly and apoptosis than did the hyperactive p53 mouse model in Mfossa et al., (2020). This is likely due to differing duration and/or intensity of p53 activation, which determines whether p53 triggers apoptosis or cell cycle arrest (Kracikova et al., 2013). The short activation of p53 induced by a single 1 Gy dose of radiation resulted in cell cycle arrest for 6 hours, premature neuronal differentiation, and transient apoptosis that spared mitotic cells lining the ventricular lumen (Mfossa et al., 2020). In contrast, permanent Aki2 loss triggered complete cell cycle exit, early differentiation, and apoptosis of not only postmitotic cells but also the neural progenitor cells that give rise to the entire cortex.

### Evidence for repression of p53 by Akirin2

Databases (Stark et al., 2006; Szklarczyk et al., 2019) as well as our own attempted co-immunoprecipitations (data not shown) provide no evidence to support a physical interaction between Aki2 and p53. While this does not eliminate the possibility that such an interaction exists, it is worth considering alternative mechanisms through which Aki2 might indirectly repress p53.

First, Aki2 could repress p53 by interacting with or modifying direct p53 regulators or signaling cascades that affect p53 protein stabilization. Mouse double minute 2 (MDM2) is an E3 ubiquitin ligase that tags p53 for proteasomal degradation to maintain low p53 protein levels in quiescent cells (Haupt et al., 1997; Honda et al., 1997; Kubbutat et al., 1997; Hafner et al., 2019). Over 300 known PTMs control p53 stabilization by interfering with p53-MDM2 binding (Hafner et al., 2019). If Aki2 represses any of the many proteins that participate in signaling cascades leading to p53 PTMs, loss of Aki2 could result in p53 activation. One potential candidate is the transcription factor, E2f1. E2f1 stabilizes p53 likely through PTMs (Kowalik et al., 1998) and is significantly upregulated in the *CaMK-Cre;Aki2*^*fl/fl*^ cortex.

Second, Aki2 could suppress p53 via NF-κB signaling. Initial studies on mammalian Aki2 and the *Drosophila* orthologue, Akirin, provide several lines of evidence that Akirin proteins interact with NF-κB transcription factor family members to activate NF-κB-dependent genes in the Imd and related mammalian immune pathways (Goto et al., 2008; Bonnay et al., 2014; Tartey et al., 2014). NF-κB has a complex, but primarily antagonistic, relationship with p53 (Ak et al., 2010). For example, p53 is a tumor suppressor gene, while NF-κB is an oncogene; activation of p53 increases neuronal death concomitant with reduced NF-κB activity, while inhibition of p53 increases neuronal survival concomitant with increased NF-κB (Plesnila et al., 2007). Because activation of one opposes the other, these two transcription factors cannot be activated within the same cell (Ak et al., 2010). Thus, it is possible that Aki2 indirectly regulates p53 through currently undefined NF-κB signaling pathways. In this vein, one can imagine that Aki2 enables transcription of NF-κB target genes in wild type cells, indirectly inhibiting p53. In the absence of Aki2, however, NF-κB-Aki2-dependent genes are silenced, resulting in increased p53 expression.

### Future directions

Before closing, it should be noted that in the CaMK-Cre mutant model used here, only a subpopulation (albeit a large one) of neurons in the cortex lack Aki2. Because of this, the power of our RNASeq analysis is likely reduced, as the transcriptomes analyzed include many neurons retaining Aki2 expression. Thus, the number of DEGs and the degree of their differential expression detected here is likely an underestimate. Future studies investigating a homogeneous population of Aki2-null neurons would aid in identifying further Aki2-dependent genes, and would be a prerequisite for utilizing chromatin immunoprecipitation (ChIP)-sequencing or ATAC-Seq approaches to identify neuronal genes directly regulated by Aki2 (presumably in a complex with transcription factors and/or chromatin remodeling machinery). Nevertheless, the work presented here raises many intriguing new insights into Aki2 function in the maturing brain that are worth exploring further, including the role of p53 pathway members as functional partners, and the potential involvement of Aki2 dysregulation in a variety of human neurodegenerative disorders.

## Acknowledgements

We would like to thank Dr. Maria Noterman, Dr. Andrew Pieper, Dr. Michael Dailey, Dr. Daniel Summers, and Dr. Diane Slusarski for generously contributing antibodies, reagents, and equipment used in these studies, the Carver Center for Genomics and Iowa Institute of Human Genetics for assistance with transcriptomics and library preparation, and Dr. Sarit Smolikove and members of the Weiner Laboratory for helpful discussions. The Jackson Laboratory Shared Scientific Services are supported in part by a Basic Cancer Center Core Grant from the National Cancer Institute (CA34196). This work was supported by a Major Project Grant from the University of Iowa Office of the Vice President for Research and Economic Development to J.A.W. and J.R.M., by an Accelerator Grant from the Iowa Neuroscience Institute to J.A.W., and by NIH R01 NS055272 to J.A.W.

## Author contributions

S.L.P., P.J.B., J.R.M., and J.A.W. conceived the experiments

S.L.P., P.J.B., B.J.I., M.P., and P.B. performed the experiments and collected and analyzed the data E.B., M.P., and P.B. performed bioinformatic analyses

S.L.P., P.J.B., and J.A.W. wrote the manuscript

All authors reviewed and edited the manuscript

J.R.M., J.J.M, and R.W.B. provided expertise and supervised bioinformatic analyses

## Methods

### Mouse strains

All experiments included both male and female animals and were conducted in accordance with the University of Iowa’s Institutional Animal Care and Use Committee and NIH guidelines. *Akirin2*^*fl/fl*^ conditional mutant mice were initially provided by Dr. Osamu Takeuchi, Kyoto University (Goto et al., 2008; Bosch et al., 2016) and have been maintained in the Weiner laboratory colony long-term. These were crossed to the *CaMKIIα-Cre* line T29-1 (Tsien et al., 1996)(JAX stock #005359) to excise the floxed *Aki2* allele in many postnatal forebrain excitatory cortical neurons. These are referred to as *CaMK-Cre;Aki2*^*fl/fl*^ in the manuscript and sometimes as cKO (conditional knockout) in several figures. As there were no obvious abnormalities in *CaMK-Cre;Aki2*^fl/+^, *Aki2*^*fl/fl*^, or *Aki2*^*fl/+*^ mice, all were used as controls. Similarly *Aki2*^*fl/fl*^*;Trp53+/+* and *Aki2*^*fl/fl*^*;Trp53+/-* did not differ, thus both were used as controls in relevant experiments. To assess Cre-mediated excision and monitor knockout cells, we also crossed these mice to the *Ai14-tdTomato* Cre-reporter line (JAX stock #007914). The early telencephalic-restricted mutant *Emx1-Cre;Aki2*^*fl/fl*^ mice were detailed in Bosch et al., 2016. *Trp53* constitutive knockout mice (Jacks et al., 1994) (JAX stock #002101), *Ai14-tdTomato* reporter mice, and line T29-1 were obtained from The Jackson Laboratory (Bar Harbor, ME). All lines were maintained on a C57BL/6J background.

### Western blots

To prevent contamination of Aki2 signal from the highly-expressing blood leukocytes, mice were transcardially perfused with 15-25 ml of 1x phosphate-buffered saline (PBS). The cortex was homogenized in 1 ml RIPA buffer (0.1% SDS, 0.25% sodium deoxycholate, 1% NP-40, 0.15M NaCl, 50mM Tris-HCl (pH 7.4), 5mM NaF) containing both protease and phosphatase inhibitors (Roche) using a Dounce homogenizer and Wheaton overhead stirrer. After incubating at 4°C for 45 mins, samples were centrifuged at 16,000 X g for 10 minutes at 4°C. Protein concentrations were measured using the Pierce BCA Protein Assay Kit (Thermo Fisher Scientific). Equal amounts of proteins were separated via SDS/PAGE using TGX precast gels (Bio-Rad), transferred to nitrocellulose membranes using a TransBlot Turbo System (Bio-Rad), and analyzed using standard quantitative western blot methods (Pillai-Kastoori et al., 2020). Chloroquine treated and nontreated HeLa cells (Cell Signaling Technology #11972) were used as positive and negative controls, respectively, for LC3 probe. Cytochrome C treated or untreated Jurkat cells (Cell Signaling Technology #9663) were used as positive and negative controls, respectively, for cleaved caspase 3 probe. Membranes were blocked in 5% nonfat milk in Tris-buffered saline with 0.1% Tween-20 (TBST). Primary antibodies were diluted in 5% BSA in TBST and membranes incubated overnight at 4°C. HRP-conjugated secondary antibodies were diluted in 5% milk in TBST and incubated at RT for 1 hour. Signal was detected using SuperSignal West Pico or Femto Enhanced Chemiluminescent Substrates (Thermo Fisher Scientific) on a LI-COR Odyssey Fc Imaging system. For all quantified probes, at least 3 biological replicates (i.e., samples from 3 mice) and at least 2 technical replicates (i.e. same samples run on different gels) per genotype or condition were run on the same gel and measured from the same image. Signal was measured using Image Studio Ver5.2 (LI-COR). All targets were normalized to a housekeeping protein (β−tubulin or GAPDH) as an internal loading control. Relative protein levels were calculated as % of 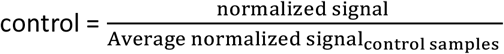. The % of control was averaged for all replicates and graphed. In instances in which protein level was measured across time, the earliest time point was used as the control in the equation above to show relative changes as maturation proceeded.

### RNA extraction and quantitative RT-PCR (qPCR)

To eliminate signal from the blood, mice older than P7 were transcardially perfused with 15-20 ml cold DEPC-treated PBS. Cortices were homogenized in 1-2 ml of TRIzol (Thermo Fisher Scientific). RNA was extracted following the manufacturer’s protocol and purified using the *mir*Vana miRNA Isolation Kit (Ambion/Life Technologies).

RNA was reverse transcribed using the High Capacity RNA-to-cDNA kit and qPCR performed using a Roche LightCycler480 with SYBR Green I Master Mix (Roche) and the primers listed in Table S2. PCR cycling parameters for 30 cycles were: 95°C 1 min, 55°C 15s, 72°C 1 min.

### Immunofluorescence

Mice were transcardially perfused with 15-20 ml of PBS followed by 15-20 ml of 4% paraformaldehyde (PFA). Brains were collected and post-fixed in 4% PFA for 2 hours – overnight at 4°C and cryoprotected in 30% sucrose. For tissues to be stained with anti-Cre antibody, 2% PFA was used for transcardial perfusion and tissue was collected directly into 30% sucrose without post-fixation. Cryoprotected tissue was quickly frozen in OCT compound (Sakura Finetek) and 18 μm coronal cryosections were cut on a Leica CM1850 cryostat. Six coronal sections per slide were collected directly onto ten slides starting at the emergence of the third ventricle and guided by reference to an atlas, ensuring that similar brain regions were compared for every mouse. Tissue was blocked for 1 hour at room temperature (RT) in PBS containing 2.5% BSA 0.5% Triton-X100 and incubated in primary antibody (Table S1) overnight at 4°C. Sections were washed in PBS and incubated in the relevant secondary antibody conjugated to Alexa-Fluor 488nm, 568nm, or 647nm (Molecular Probes/Invitrogen) for 1 hour at RT. After washing in PBS, sections were counter stained with DAPI (4’,6-diamidino-2-pheylindole) and mounted in Fluoro-Gel (Electron Microscopy Services #17985-11).

#### *In situ* hybridization

Digoxigenin-UTP (Roche)-labeled *Aki2* sense and antisense riboprobes were generated per manufacturer’s protocols as in (Bosch et al., 2016). *In situ* hybridization was performed following previously published methods (Grove et al., 1998; Wang et al., 2002). Fresh-frozen tissue was cryosectioned at 20 µm and dried at 65°C for 10 mins followed by RT for 10 mins. Tissue was fixed in 4% PFA for 10 min at RT followed by washes in PBS and a 10 min incubation in fresh acetylation solution (0.1M triethanolamine, 0.65% HCl, 0.25% acetic anhydride). After washing in PBS, slides were incubated in hybridization solution (50% formamide, 5x SSC, 5x Denhart’s solution, 250μg/ml tRNA, 500μg/ml salmon sperm DNA, 50μg/ml heparin) at RT for 2 hours. Slides were incubated with riboprobes at 65°C overnight. The following day, slides were washed over 4 hours in 0.2x SSC at 65°C followed by washes in TBST at RT. Slides were blocked in blocking solution (Roche DIG wash and block buffer set) diluted in 1x maleic acid for 1h at RT and riboprobes were detected by incubating in anti-digoxigenin-AP antibody (1:2000 in 1% sheep serum, Roche) overnight at 4°C. After several washes in TBST over 15 mins, slides were incubated in alkaline buffer for 10 mins (100mM NaCl, 100mM Tris-HCl (pH 9.5), 50mM MgCl_2_, 1% Tween-20), and developed using nitro blue tetrazolium (NBT) and 5-bromo-4-chloro-3-indolyl phosphate (BCIP) at RT until sufficient color developed.

### Imaging

A Leica SPE TCS Confocal Microscope and Leica Application Suite software were used for confocal and epifluorescent imaging. All epifluorescent and confocal images were captured using the same settings for each antibody. *In situ* hybridization was imaged on a Zeiss Axiophot microscope equipped with an Axiocam Hrc and using AxioVision Rel4.8. Images of silver and Golgi staining were captured using a Leica DMIRB inverted microscope. Brightness and contrast were adjusted equally on images using ImageJ/FIJI (Schindelin et al., 2012; Schneider et al., 2012) or Adobe Photoshop.

### Cell counts

For cell counts, 3-6 coronal sections per mouse were imaged at the convergence of the primary motor and somatosensory cortices and analyzed using FIJI. Only brain regions determined to be similar based on histological landmarks and with reference to an atlas were included for analysis.

Cux1-positive cells in layers II-IV and FoxP2-positive cells in layers VI were counted in one (10x) field of view per section using the FIJI cell counter plug-in. Ctip2 expression, cell size, and cell density were used to identify Layer V, in which cells were counted in the same manner.

For NeuN, GFAP, and CD68 measurements, two 10x fields of view vertically spanning the cortex from white matter to pia were montaged in Adobe Photoshop. The background was subtracted with a rolling ball radius of 12 pixels and NeuN-positive cells were counted using analyze particles plugin with size = 20.5 – infinity; circularity = 0.2-1.00 in FIJI. The resulting images were manually inspected for mis-identified or conjoined cells for which the numbers were properly adjusted. On the same images, threshold was set to a level that optimized signal while reducing background. The individual areas of 10 cells in each of layers II-III, IV, V, and VI were measured in each image using FIJI. The average cell area in each layer per section is reported from at least 3 mice per genotype per age.

Average fluorescence intensity of GFAP staining and the area of CD68 as a percentage of the entire cortical area was measured within a polygon covering the entire cortex including white matter but excluding subcortical structures. The length of a line drawn from the dorsal edge of the lateral ventricle (including the white matter) to the dorsal edge of the pia (including Layer I) was used to estimate cortical thickness.

### NeuroSilver and Golgi staining

For NeuroSilver staining, animals were transcardially perfused as for immunofluorescence, but replacing 1x PBS with 0.1M phosphate buffer (pH 7.4). Sections were cut on a Leica CM1850 cryostat at 60 µm and incubated in 4% PFA for 7 days at 4°C. Tissue was stained using the FD NeuroSilver Kit II (FD Neurotechnologies, Inc. Cat #PK301A) according to manufacturer’s protocol. For Golgi staining, the two cerebral hemispheres were separated from the rest of the brain and incubated with the solutions of the FD Rapid GolgiStain Kit (FD NeuroTechnologies, Inc) according to manufacturer’s protocol. One hundred-µm cryostat sections of telencephalon were mounted on slides and examined directly.

### TUNEL staining

Eighteen-µm cryostat sections from the cerebrum were fixed in 4% PFA directly on slides and stained using the FragEL DNA Fragmentation Kit (QIA33; Millipore Sigma) according to manufacturer’s protocol. A positive control was generated by treating cortical sections with 20 μg/ml proteinase K for 10 minutes at RT followed by 1μg/μl DNase I in 1xTBS/1mM MgSO4 for 20 minutes at RT.

### Nec-1s administration

Nec-1s (7-Cl-O-Nec1; Abcam #ab221984) was dissolved in DMSO, diluted in sterile saline, and administered at 6.25mg/kg via intraperitoneal injections daily beginning at P21 and ending at P50. Three to four mice per genotype were injected with Nec-1s or the equivalent amount of DMSO vehicle.

### Statistics

All statistical tests were performed using GraphPad Prism Software with significance level p<0.05. Outlier data points, determined using the ROUT method with a Q=1, were excluded.

### Transcriptomic analyses

#### *CaMK-Cre;Aki2*^*fl/fl*^ transcriptome

After transcardial perfusion with 15-20ml of cold DEPC-treated PBS, total RNA was extracted as above from the cortices of 3 *Aki2*^*wt/wt*^ and 1 *CaMK-Cre;Aki2*^*wt/wt*^ controls and 3 *CaMK-Cre;Ak2*^*fl/fl*^ mutants at P35. mRNA-enriched libraries were prepared by the Carver Center for Genomics (University of Iowa) using the Illumina TruSeq Stranded mRNA Library Prep kit and quantified using Turner BioSystems TBS-380 fluorometer and Invitrogen Quant-It Picogreen. The Genome Technologies service at The Jackson Laboratory (Bar Harbor, ME) subsequently QC’ed the libraries on an Agilent Bioanalyzer and sequenced them on an Illumina NextSeq 500, with 75 bp single end reads. At least 49M reads per sample were generated.

RNA used for qPCR validation was treated with DNase using the TURBO DNA-free Kit (Thermo Fisher) according to manufacturer’s protocol. Concentration and quality were assessed using a Trinean DropSense-16 Spectrophotometer and Agilent Bioanalyzer. All samples used had an RIN of 8.6 or higher. Two µg of RNA were reverse transcribed using the High-Capacity RNA-to-cDNA Kit (Applied Biosystems/Thermo Fisher). qPCR was performed as above. The primers used are listed in Table S2. All qPCR validating CaMK-Cre transcriptomics used ribosomal protein L32 (RPL32) and transferrin receptor (Tfrc) as reference genes and relative expression levels were quantified using qbase + software (Biogazelle, Zwijnaarde, Belgium www.qbaseplus.com), employing the method described by (Vandesompele et al., 2002). Statistical differences were determined using two-tailed unpaired t-tests on log-transformed CNRQ values.

#### *Emx1-Cre;Aki2*^*fl/fl*^ transcriptome

The telencephalon was carefully dissected from E10.5 embryos in cold PBS and placed into RNAlater (Invitrogen) at 4°C for a maximum of 1 week. RNAlater was carefully removed and the RNA was extracted and purified as above. RNA samples were assessed for purity using a nanodrop and for RNA integrity using the Experion automated electrophoresis station (BioRad). All samples used in the RNAseq experiments had an RQI of >8. Library preparation was performed using the Illumina TruSeq Stranded mRNA library prep kit and RNA sequencing was performed at the Iowa Institute of Human Genetics (University of Iowa). Libraries were generated with unique barcodes, combined equally into one pool and sequenced over 2 lanes of the Illumina HiSeq4000, using 75bp paired end read lengths.

For qPCR validation studies, RNA (90-100ng) collected from E10.5 telencephalon was reverse transcribed using the High Capacity RNA-to-cDNA kit and qPCR performed as above using the primers listed in Table S2. The ddC(t) method was used to calculate fold change using β-Actin as a reference gene as described previously (Bosch et al., 2019).

### Bioinformatics

RNASeq data were processed with the bcbio-nextgen pipeline (https://github.com/bcbio/bcbio-nextgen, version 1.1.4). The pipeline uses STAR (Dobin et al., 2013) to align reads to the mm10 genome build (GENCODE release M10, Ensembl 89 annotation) and quantifies expression at the gene level with featureCounts (Liao et al., 2014). All further differential expression analyses were performed using R (R Core Team, 2019). For gene level count data, the R package EDASeq (Risso et al., 2011) was used to adjust for GC content effects (full quantile normalization) and account for sequencing depth (upper quartile normalization). Latent sources of variation in expression levels were assessed and accounted for using RUVSeq (RUVr mode using all features) (Risso et al., 2014). Differential expression analysis was conducted using the edgeR quasi-likelihood pipeline (Robinson et al., 2010; McCarthy et al., 2012; Chen et al., 2016). Gene ontology and KEGG pathway analysis was performed using the R package clusterProfiler (Yu et al., 2012). Ingenuity Pathway Analysis (Qiagen Redwood City, CA, USA, www.qiagen.com/ingenuity) (Krämer et al., 2013) was used for upstream analysis with molecule type restricted to transcriptional regulator.

All RNASeq data have been deposited in the NCBI Gene Expression Omnibus database, accession number GSE178844 (https://www.ncbi.nlm.nih.gov/geo/query/acc.cgi?acc=GSE178844).

