## Supplemental Files for "p53-mediated neurodegeneration in the absence of the nuclear protein Akirin2"

### Supplemental information:

**Supplemental Figure Legends**

**Supplemental Figure 1: Akirin2 transcript expression and CaMKIIα-Cre (line T29-1) activity in the postnatal brain**

A) *In situ* hybridization of adult mouse cerebral cortex demonstrates ubiquitous expression of Aki2 transcript (antisense signal, with sense negative control) across all layers (I-VI), the white matter (wm), and hippocampus (hipp). B) Diffusion of tdTomato protein into neuropil obfuscate individual Cre-expressing cell bodies by P35, C) while earlier imaging of the sagittal cortex shows Cre expression mostly limited to hippocampal CA1 and cortical layers II-IV and VI rostral (R) to the visual cortex. D, dorsal; V, ventral; R, rostral; C, caudal; olf, olfactory bulb; LV, lateral ventricle. Scale bar: A) 100 μm B,C) 1500 μm.

**Supplemental Figure 2: Cell-autonomous loss of FoxP2-negative layer VI neurons in Akirin2 mutant cortex**

A tdTomato Cre reporter (Figure 1E) indicates that FoxP2-negative neurons are Cre-positive. Cell quantification (Figure 3D) suggests that initial cell death occurs in Cre-positive cells. To confirm this, cortical cryosections from a control (n=5 sections from 1 mouse) and an Aki2 mutant (n=6 sections from 1 mouse) at P50 were immunostained with antibodies against FoxP2 and NeuN. FoxP2-negative/NeuN-positive cells were quantified. As graphed at left, a significant loss of these neurons occurs in Aki2 mutants. Data shown as mean ± SEM. $**p<0.01$ determined using unpaired two-tailed t-test. Scale bar, 200 μm.

**Supplemental Figure 3. Lack of evidence for apoptosis or autophagy in the Akirin2 mutant cortex**

Neither immunostaining (A) nor western blotting (B) of cortical tissues for the apoptotic marker cleaved caspase-3 (CC3) indicates any increase in apoptosis in *CaMK-Cre;Aki2*$^{fl/fl}$ mutants (cKO) at P28, P35, or later ages (not shown; GAPDH antibody used as loading control on western blots). C) Similarly, TUNEL staining for DNA fragmentation, a hallmark of apoptotic cells, indicates little if any apoptosis in Aki2 mutant cortex cryosections at any age examined, despite clear staining of DNase-treated positive control sections. D) Double-staining of control cortical cryosections with NeuroTrace (NT)500, a fluorescent Nissl stain, and an antibody against NeuN. The two signals overlap nearly completely (graph shows near-perfect correlation between NeuN and NT500 cell area), confirming the NeuN labels neuronal somatic cytoplasm as well as nuclei (n=79 cells from 6 cryosections in 1 mouse). E) Western blot of cortical lysates from control and Aki2 mutants at the indicated ages probed with an antibody against LC3, indicates no sign of increased autophagy in the absence of Aki2. Positive control samples (chloroquine treated HeLa cells) show the expected increase in lipidated LC3 II band, which runs at a lower apparent molecular weight compared to the non-lipidated LC3 I band. β-tubulin (βtub) antibody used as loading control. Correlation determined using Pearson's r. Scale bar = 200μm.

**Supplemental Figure 4. Transcriptomic analysis of Akirin2 mutant embryonic telencephalon.**

RNA extracted from E10.5 *Emx1-Cre;Aki2$^{fl/fl}$* and control telencephalon was sequenced and bioinformatics was performed as described in Materials and Methods. A) Volcano plot of log FC as a function of –log(10) FDR of all identified genes shows 295 genes significantly upregulated (red) and 156 genes significantly downregulated (blue) in *Emx1-Cre;Aki2$^{fl/fl}$* mutants (n=4) compared to controls (n=4) with an FDR cut off of 0.05. Genes involved in p53 pathway are highlighted in black. Note significant decrease in Aki2 highlighted in purple. B-C) Gene ontology (GO) analysis bar plot showing significant biological processes enriched in the list of genes with FDR ≤ 0.05 (B) as well as FDR ≤ 0.1 (C). Top 20 significant biological processes are shown in GO bar blot with bar color corresponding to p-value as delineated in legend. D) Gene-concept network showing the relationship of several dysregulated genes to the highlighted GO terms in (B). Size of black circles associated with GO terms indicates the number of genes involved while color of gene circles indicates signed -log10(FDR) as indicated in the legend. E) qPCR validation of several p53 target genes on control (n=6-8) and *Emx1-Cre;Aki2$^{fl/fl}$* (n=4-8) biological replicates. F) KEGG pathway analysis of DEGs in both embryonic transcriptome (orange) and postmitotic transcriptome (green) shows transcriptional dysregulation converges on the p53 signaling pathway. The size of the circle associated with the pathway indicates the number of genes involved as delineated in the legend. Data shown as mean ± SEM. **p<0.01, ***p<0.001 as determined by unpaired two-tailed t-test.

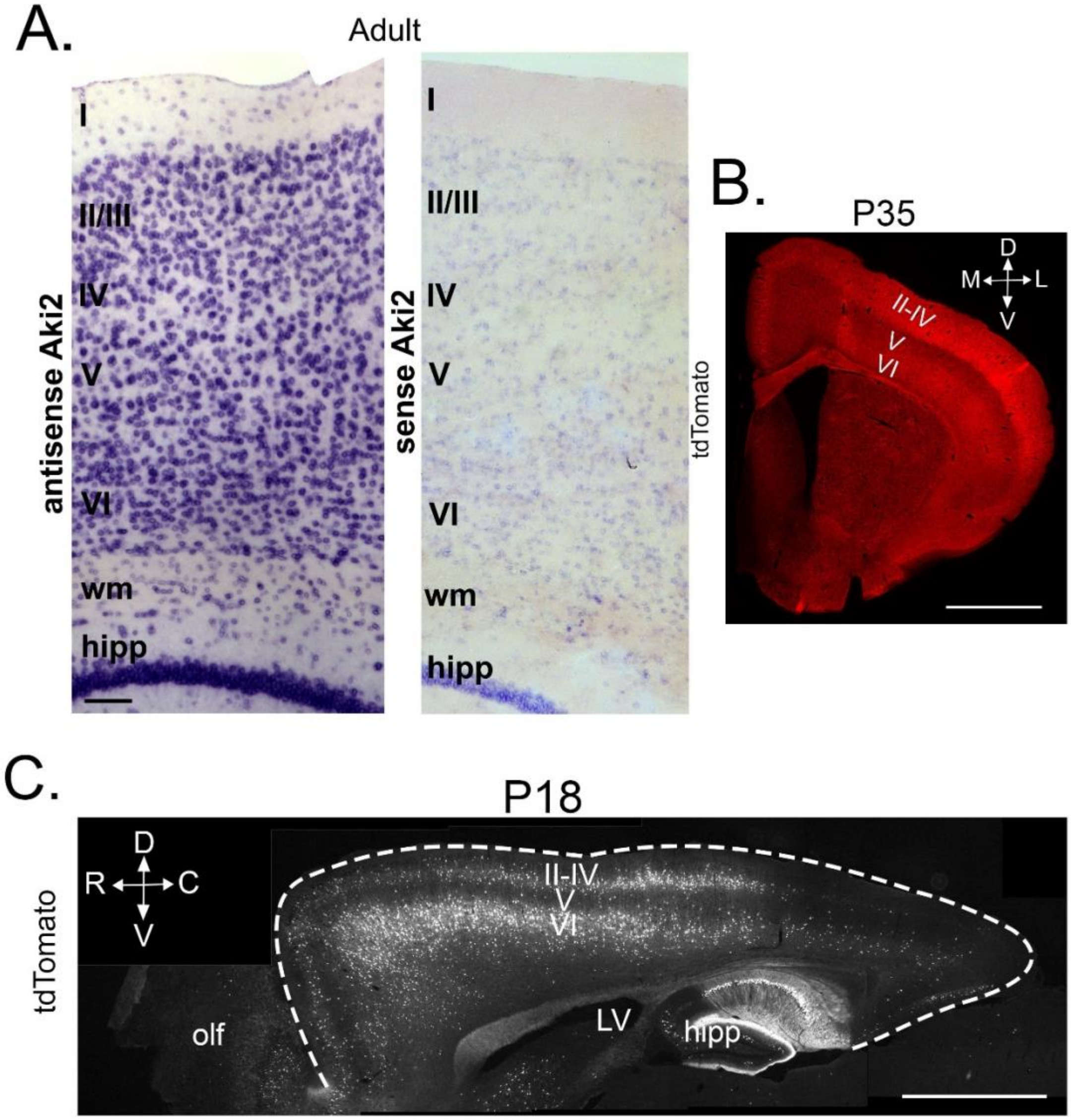


**Supplemental Figure 1**

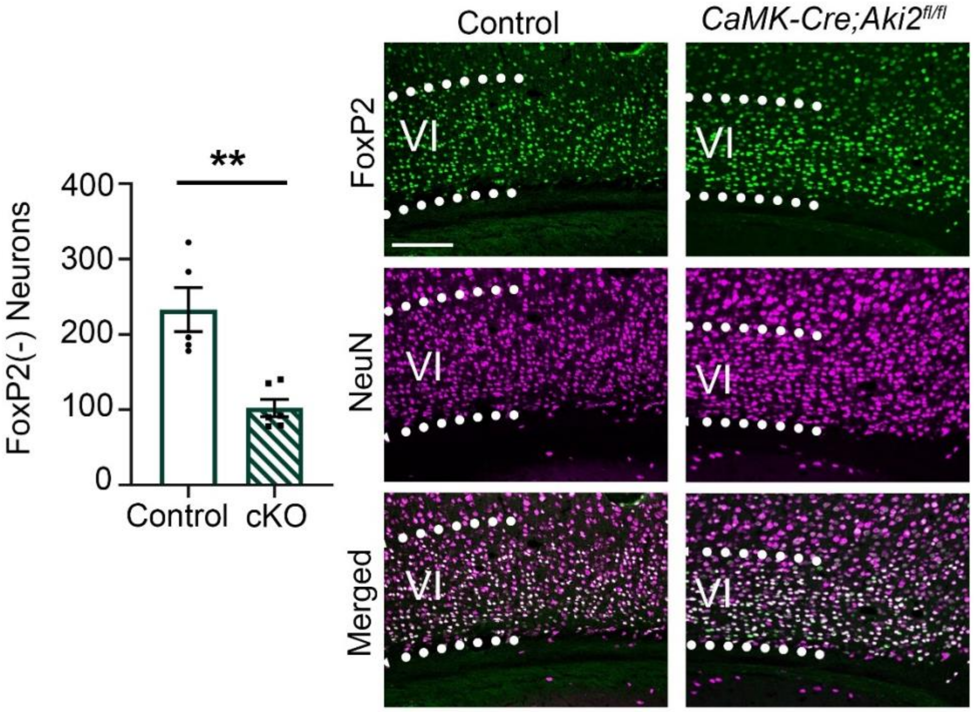


**Supplemental Figure 2**

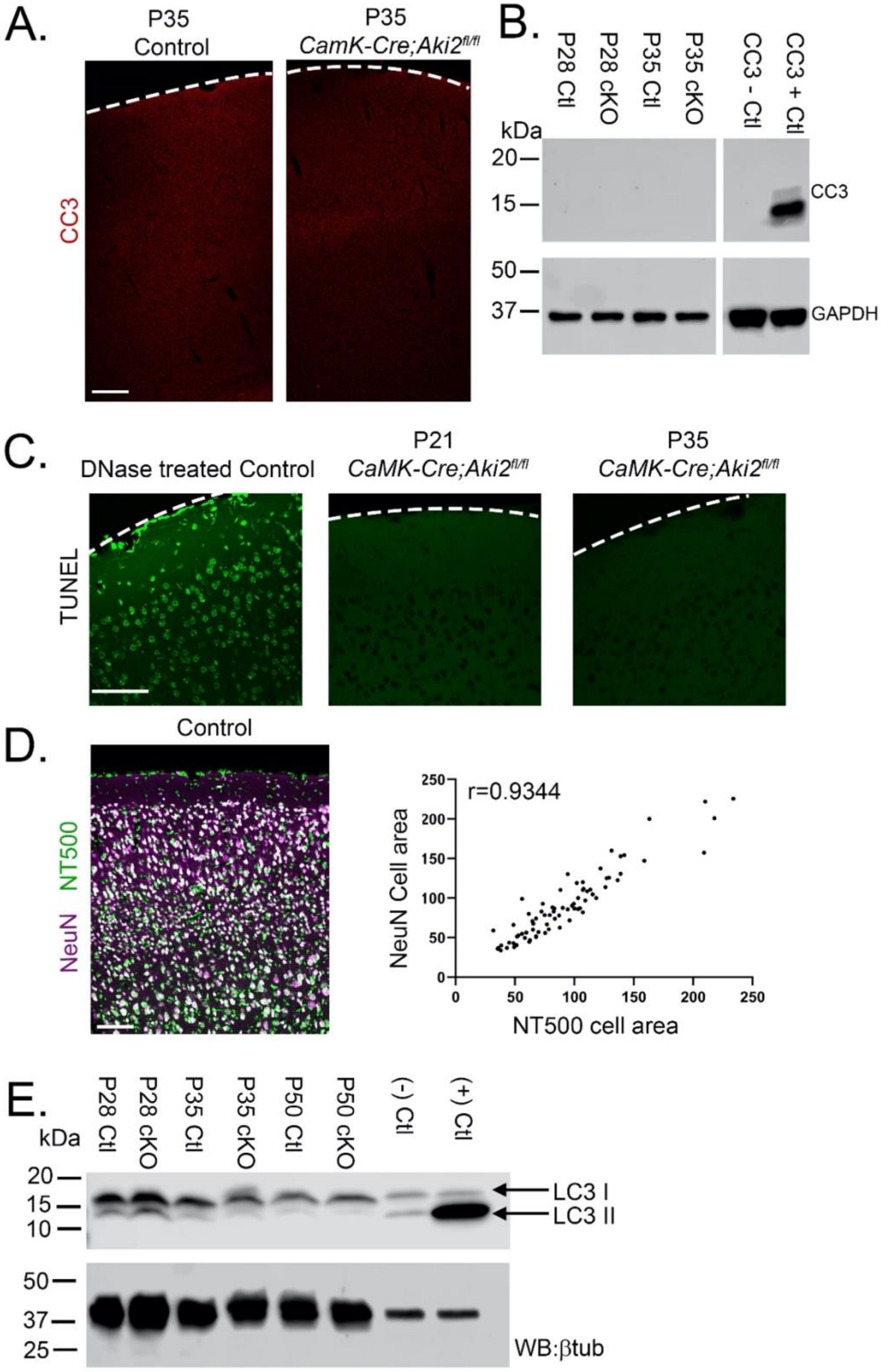


**Supplemental Figure 3**

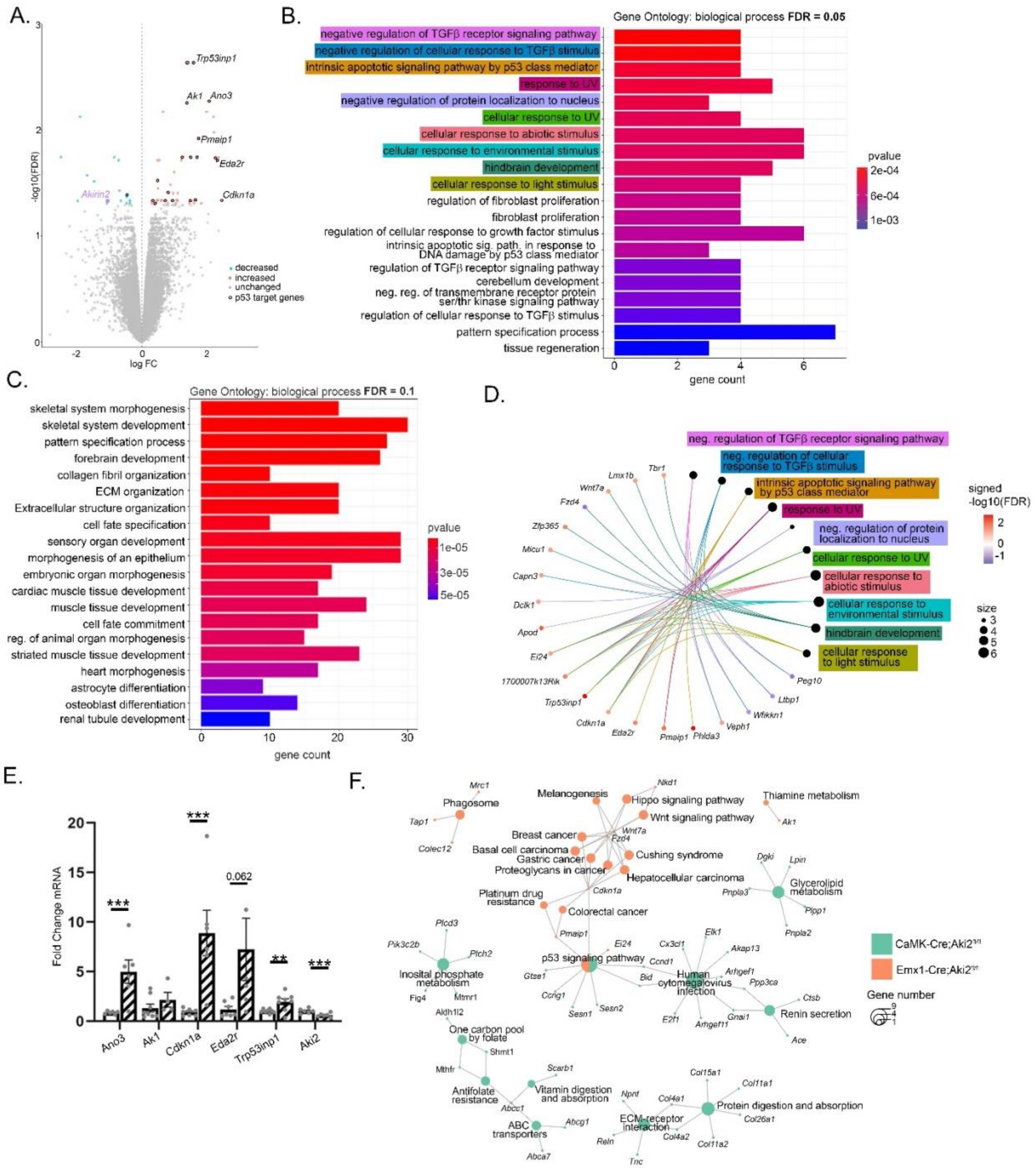


**Supplemental Figure 4**

| Antibody | IF Dilution | WB dilution | Source | Identifier |
|---|---|---|---|---|
| **Akirin2** | 1:500 | 1:1000 | Abcam | Ab221475 |
| **β–tubulin** | | 1:1000 | Sigma-Aldrich | T4026 |
| **Cleaved-caspase-3 (CC3)** | 1:100-1:500 | 1:1000 | Cell Signaling Technologies | 9661 |
| **CD68** | 1:500 | | Bio-Rad | MCA1957T |
| **Cre** | 1:500 | | Millipore | Mab3120 |
| **Ctip2** | 1:200 | | Abcam | Ab18465 |
| **Cux1** | 1:100 | | Santa Cruz Biotechnologies | SC 13024 |
| **FoxP2** | 1:500 | | Abcam | Ab16046 |
| **GAPDH** | | 1:1000 | Abcam | Ab8245 |
| **GFAP** | 1:500 | | Sigma-Aldrich | G3893 |
| **NeuN** | 1:300 | | Millipore | Mab377 |
| **LC3B** | | 1:1000 | Cell Signaling Technologies | 2775 |
| **p53/Trp53 (1C12)** | 1:500 | 1:1000 | Cell Signaling Technologies | mAb #2524 |
| **P2Y12** | 1:1000 | | AnaSpec | AS-55043A |
| **RIPK1 (D94C12)** | | 1:1000 | Cell Signaling Technologies | 3493 |
| **RIPK3 (D8JSL)** | | 1:1000 | Cell Signaling Technologies | 15828 |

Supplemental Table 1: Primary antibodies employed

| Gene | Forward Primer | Reverse Primer |
|---|---|---|
| **Aki2** | CTTCGGCGACGTCTCTTCC | ATGAAGTCCCTGGTGATGCT |
| **Ak1** | TCCACCTCCATGTAAAGGCTG | AAGCAGGTCCCCAGTAGACA |
| **Ano3** | AAGGACTCTACCCTTAAGTGTTCC | GGAGACGAGATCGATCATAGTTGG |
| **Bid** | TCTGAGGTCAGCAACGGTTC | CTCTTGGCGAGTACAGCCAG |
| **Cdkn1a** | TGAGGAGGAGCATGAATGGAG | GGAACAGGTCGGACATCACC |
| **Dbf4** | AATAAGATACAGTGTCGGGTCCC | GTCCTTCTGGAAATTGGGCTC |
| **Eda2r** | CCTACCTCAGATGGGACATCA | CGGTACTCATTCTCTTGACAATCCA |
| **Gmnc** | TCTGGAAGAGAAGGCCAAGA | CCCAGGTTGTTCCTCACAGT |
| **Gtse1** | TTTTGGGCCTGTTGGACATAAA | CTCAAGGTGCAAGGGCTACC |
| **Met** | GTGAACATGAAGTATCAGCTCCC | TGTAGTTTGTGGCTCCGAGAT |
| **Pls3** | GCGACCACCCAGATTTCCAAA | GCAGTGGCATATTAGCTTCCTTG |
| **RPL32** | AGTTCATCAGGCACCAGTCA | GCTCCTTGACATTGTGGACC |
| **Tex15** | CTCCAGGTACAGAGTGCACAG | AGGAGATCCTGTAGTGCTTGGA |
| **Tfrc** | GATCAAGCCAGATCAGCATTCT | ACCGGGTGTATGACAATGGTT |
| **Trp53** | CCCAGCGAAATTCTATCCAG | CAGACAGGCTTTGCAGAATG |
| **Trp53inp1** | GTGTGCTCTGCTGAGGACTC | GTTGACTTCATAGATACCTGCCC |
| **Ttc28** | GGTACTGCCTATCGAATGGTCC | GCTCCTGGTAACACTTGAACG |
| **β-Actin** | CACTGTCGAGTCGCGTCC | TCATCCATGGCGAACTGGTG |

Supplemental Table 2: qPCR primer sequences

| | | Emx1 Transcriptome | | | CaMK Transcriptome | | | |
|---|---|---|---|---|---|---|---|---|
| **Symbol** | **Gene Name** | **Log FC** | **P value** | **FDR** | **Log FC** | **P value** | **FDR** | **direction** |
| Trp53inp 1 | transformation related protein 53 inducible nuclear protein 1 | 1.60 | 3.16E-07 | 0.0023 2 | 0.371 | 2.51E -04 | 0.027 | up in both |
| CCng1 | cyclin G1 | 1.42 | 4.27E-07 | 0.0023 2 | 0.260 | 5.04E -04 | 0.03495 | up in both |
| Espn | espin | -2.50 | 1.67E-05 | 0.0180 7 | -0.962 | 6.78E -05 | 0.01683 5 | down in both |
| Eda2r | ectodysplasin A2 receptor | 2.34 | 2.26E-05 | 0.0193 1 | 1.207 | 5.39E -04 | 0.03545 8 | up in both |
| Susd6 | sushi domain containing 6 | 0.54 | 2.68E-05 | 0.0193 1 | 0.228 | 7.07E -04 | 0.03862 4 | up in both |
| Pls3 | plastin 3 (T-isoform) | 0.31 | 0.00019 6 | 0.0492 0 | 0.319 | 3.61E -04 | 0.03131 1 | up in both |

Supplemental Table 3: Differentially expressed genes common to both embryonic cortical progenitor and postmitotic neuron transcriptomes

| Upstream Regulator | Expr Log Ratio | Molecule Type | Predicted Activation State | Activation z-score | p-value of overlap |
|---|---|---|---|---|---|
| TP53 | 0.228 | transcription regulator | Activated | 2.27 | 9.82E-05 |
| TP73 | -0.091 | transcription regulator | | 0.644 | 3.11E-04 |
| FOXO3 | -0.11 | transcription regulator | | 1.849 | 3.22E-04 |
| MYCN | -0.086 | transcription regulator | | -0.956 | 5.12E-04 |
| SOX21 | -0.2 | transcription regulator | | | 8.84E-04 |
| CTNNB1 | -0.015 | transcription regulator | | 0.096 | 1.15E-03 |
| EIF2S1 | -0.053 | translation regulator | | | 2.38E-03 |
| SOX17 | 0.048 | transcription regulator | | 1.941 | 2.71E-03 |
| KMT2D | 0.187 | transcription regulator | | 1.353 | 2.72E-03 |
| GLI1 | 0.286 | transcription regulator | | -0.576 | 3.84E-03 |
| PAX6 | -0.018 | transcription regulator | | 0.365 | 3.84E-03 |
| TWIST2 | -0.029 | transcription regulator | | 0.068 | 4.08E-03 |
| CDKN2A | | transcription regulator | | -0.107 | 5.83E-03 |
| ISX | | transcription regulator | | | 5.91E-03 |
| MED1 | 0.083 | transcription regulator | | 0.314 | 6.57E-03 |
| NANOG | | transcription regulator | | 0.447 | 7.23E-03 |
| ID1 | -0.18 | transcription regulator | | 0.115 | 7.71E-03 |
| EMX2 | -0.116 | transcription regulator | | | 7.79E-03 |
| MNX1 | | transcription regulator | | | 9.91E-03 |
| Fus | -0.288 | transcription regulator | | | 9.93E-03 |
| EP300 | 0.027 | transcription regulator | | -0.351 | 1.03E-02 |
| TP63 | 0.242 | transcription regulator | | 0.827 | 1.04E-02 |
| RB1 | 0.024 | transcription regulator | Inhibited | -2.37 | 1.04E-02 |
| FOXM1 | -0.23 | transcription regulator | | 0.316 | 1.07E-02 |
| FOXO4 | -0.044 | transcription regulator | | 0.816 | 1.22E-02 |

Supplemental Table 4: Top 25 predicted upstream regulators of the *CaMK-Cre;Aki2*$^{fl/fl}$ transcriptome

| Upstream Regulator | Expr Log Ratio | Molecule Type | Predicted Activation State | Activation z-score | p-value of overlap |
|---|---|---|---|---|---|
| TP53 | -0.148 | transcription regulator | Activated | 3.503 | 4.92E-10 |
| MDM2 | 0.275 | transcription regulator | | | 0.00000375 |
| CTNNB1 | 0.112 | transcription regulator | | 1.854 | 0.00000827 |
| GATA2 | -1.226 | transcription regulator | Inhibited | -2.138 | 0.0000164 |
| ASCL1 | 0.6 | transcription regulator | | 1.067 | 0.000127 |
| KMT2A | -0.23 | transcription regulator | | | 0.000127 |
| IGF1 | -0.09 | growth factor | | 0.603 | 0.000201 |
| HDAC8 | 0.061 | transcription regulator | | | 0.000325 |
| PITX2 | | transcription regulator | | | 0.000453 |
| WBP2 | -0.08 | transcription regulator | | 0 | 0.000522 |
| MYF5 | | transcription regulator | | | 0.000652 |
| NDN | 0.102 | transcription regulator | | | 0.000856 |
| TWIST1 | -0.467 | transcription regulator | | -1.206 | 0.00112 |
| CDKN2A | | transcription regulator | | 1.412 | 0.00129 |
| KLF6 | 0.378 | transcription regulator | | | 0.00137 |
| BMP7 | 0.32 | growth factor | | 0.975 | 0.00148 |
| BRCA1 | -0.141 | transcription regulator | | 1.976 | 0.00148 |
| GDF15 | | growth factor | | | 0.00149 |
| HDAC7 | -0.128 | transcription regulator | | | 0.00149 |
| HMGB1 | | transcription regulator | | | 0.00168 |
| BRD7 | -0.098 | transcription regulator | | | 0.00178 |
| TGFB1I1 | 0.164 | transcription regulator | | | 0.00194 |
| PHF1 | 0.107 | transcription regulator | | | 0.00228 |
| JUN | 0.177 | transcription regulator | | | 0.00235 |
| ESRRB | 0.346 | transcription regulator | | | 0.00246 |

Supplemental Table 5: Top 25 predicted upstream regulators of the *Emx1-Cre:Aki2*$^{fl/fl}$ transcriptome

| ID | Genes in dataset | Expr Log Ratio | Direction of p53 regulation | Source |
|---|---|---|---|---|
| ENSMUSG00000023088 | ABCC1 | 0.408 | Downregulates | Ingenuity Knowledge Base |
| ENSMUSG00000020681 | ACE | 0.925 | Regulates | Ingenuity Knowledge Base |
| ENSMUSG00000018340 | ANXA6 | 0.279 | Upregulates | Ingenuity Knowledge Base |
| ENSMUSG00000025533 | ASL | 0.555 | Upregulates | Ingenuity Knowledge Base |
| ENSMUSG00000016252 | Atp5e | -0.223 | Downregulates | Ingenuity Knowledge Base |
| ENSMUSG00000030268 | BCAT1 | 0.205 | Regulates | Ingenuity Knowledge Base |
| **ENSMUSG00000004446** | **BID** | **0.491** | **Upregulates** | **Ingenuity Knowledge Base** |
| ENSMUSG00000022098 | BMP1 | -0.317 | Upregulates | Ingenuity Knowledge Base |
| ENSMUSG00000022665 | CCDC80 | 0.344 | Upregulates | Ingenuity Knowledge Base |
| ENSMUSG00000070348 | CCND1 | -0.371 | Regulates | Ingenuity Knowledge Base |
| ENSMUSG00000020326 | CCNG1 | 0.26 | Upregulates | Ingenuity Knowledge Base |
| ENSMUSG00000023473 | CELSR3 | 0.444 | Downregulates | Ingenuity Knowledge Base |
| ENSMUSG00000020741 | CLUH | 0.212 | Regulates | Ingenuity Knowledge Base |
| ENSMUSG00000031502 | COL4A1 | 0.727 | Upregulates | Ingenuity Knowledge Base |
| ENSMUSG00000031503 | COL4A2 | 0.566 | Downregulates | Ingenuity Knowledge Base |
| ENSMUSG00000028247 | COQ3 | -0.348 | Downregulates | Ingenuity Knowledge Base |
| ENSMUSG00000021939 | CTSB | 0.279 | Downregulates | Ingenuity Knowledge Base |
| ENSMUSG00000031778 | CX3CL1 | 0.579 | Upregulates | Ingenuity Knowledge Base |
| **ENSMUSG00000002297** | **DBF4** | **0.668** | **Downregulates** | **Ingenuity Knowledge Base** |
| ENSMUSG00000028035 | DNAJB4 | 0.306 | Downregulates | Ingenuity Knowledge Base |
| ENSMUSG00000034457 | EDA2R | | Upregulates | Ingenuity Knowledge Base |
| ENSMUSG00000027490 | E2F1 | 0.391 | Regulates | Ingenuity Knowledge Base |
| ENSMUSG00000064080 | FBLN2 | 0.34 | Upregulates | Ingenuity Knowledge Base |
| ENSMUSG00000020635 | FKBP1B | 0.534 | Upregulates | Ingenuity Knowledge Base |
| ENSMUSG00000097248 | Gm2694 | 0.553 | Downregulates | Ingenuity Knowledge Base |
| ENSMUSG00000057614 | GNAI1 | 0.203 | Upregulates | Ingenuity Knowledge Base |
| **ENSMUSG00000022385** | **GTSE1** | **0.928** | **Upregulates** | **Ingenuity Knowledge Base** |
| ENSMUSG00000020644 | ID2 | -0.211 | Downregulates | Ingenuity Knowledge Base |
| ENSMUSG00000028438 | KIF24 | 0.814 | Regulates | Ingenuity Knowledge Base |
| ENSMUSG00000073700 | KLHL21 | 0.359 | Regulates | Ingenuity Knowledge Base |
| ENSMUSG00000020593 | LPIN1 | 0.458 | Upregulates | Ingenuity Knowledge Base |
| ENSMUSG00000024948 | MAP4K2 | 0.225 | Downregulates | Ingenuity Knowledge Base |
| ENSMUSG00000024556 | ME2 | 0.386 | Downregulates | Ingenuity Knowledge Base |
| **ENSMUSG00000009376** | **MET** | **0.57** | **Downregulates** | **Ingenuity Knowledge Base** |
| ENSMUSG00000075705 | MSRB1 | 0.407 | Upregulates | Ingenuity Knowledge Base |
| ENSMUSG00000031762 | Mt2 | 0.5 | Upregulates | Ingenuity Knowledge Base |
| ENSMUSG00000022255 | MTDH | -0.224 | Regulates | Ingenuity Knowledge Base |
| ENSMUSG00000017774 | MYO1C | 0.459 | Upregulates | Ingenuity Knowledge Base |
| ENSMUSG00000037966 | NINJ1 | 0.363 | Upregulates | Ingenuity Knowledge Base |

| | | | | |
|---|---|---|---|---|
| ENSMUSG00000038146 | NOTCH3 | -0.321 | Regulates | Ingenuity Knowledge Base |
| ENSMUSG00000040998 | NPNT | 0.236 | Regulates | Ingenuity Knowledge Base |
| ENSMUSG00000035232 | PDK3 | 0.313 | Upregulates | Ingenuity Knowledge Base |
| ENSMUSG00000030880 | POLR3E | 0.249 | Upregulates | Ingenuity Knowledge Base |
| ENSMUSG00000028161 | PPP3CA | -0.454 | Downregulates | Ingenuity Knowledge Base |
| ENSMUSG00000027540 | PTPN1 | 0.232 | Regulates | Ingenuity Knowledge Base |
| ENSMUSG00000021091 | SERPINA3 | 0.896 | Upregulates | Ingenuity Knowledge Base |
| ENSMUSG00000038332 | SESN1 | 0.39 | Upregulates | Ingenuity Knowledge Base |
| ENSMUSG00000028893 | SESN2 | 0.418 | Upregulates | Ingenuity Knowledge Base |
| ENSMUSG00000074637 | SOX2 | -0.384 | Upregulates | Ingenuity Knowledge Base |
| ENSMUSG00000021611 | TERT | 0.827 | Downregulates | Ingenuity Knowledge Base |
| **ENSMUSG00000009628** | **TEX15** | **1.997** | **Upregulates** | **Ingenuity Knowledge Base** |
| ENSMUSG00000030095 | TMEM43 | 0.33 | Regulates | Ingenuity Knowledge Base |
| ENSMUSG00000028364 | TNC | -0.754 | Upregulates | Ingenuity Knowledge Base |
| **ENSMUSG00000028211** | **TP53INP1** | **0.371** | **Upregulates** | **Ingenuity Knowledge Base** |
| ENSMUSG00000000296 | TPD52L1 | 0.295 | Regulates | Ingenuity Knowledge Base |
| **ENSMUSG00000033209** | **TTC28** | **1.023** | **Regulates** | **Ingenuity Knowledge Base** |
| ENSMUSG00000029512 | ULK1 | 0.362 | Upregulates | Ingenuity Knowledge Base |

Supplemental Table 6: p53 target genes dysregulated in the *CaMK-Cre;Aki2*$^{fl/fl}$ transcriptome. Bolded genes were validated via qPCR.

| ID | Genes in dataset | Expr Log Ratio | Direction of p53 regulation | Source |
|---|---|---|---|---|
| **ENSMUSG00000074968** | **Ano3** | | | **Quintens et al., 2015** |
| **ENSMUSG00000023067** | **CDKN1A** | **2.472** | **Upregulates** | **Ingenuity Knowledge Base** |
| **ENSMUSG00000034457** | **EDA2R** | **2.34** | **Upregulates** | **Ingenuity Knowledge Base** |
| ENSMUSG00000026831 | C9orf116 | 2.281 | Upregulates | Ingenuity Knowledge Base |
| ENSMUSG00000048458 | INKA2 | 2.003 | Upregulates | Ingenuity Knowledge Base |
| ENSMUSG00000024521 | Pmaip1 | 1.755 | Upregulates | Ingenuity Knowledge Base |
| ENSMUSG00000046818 | DDIT4L | 1.708 | Upregulates | Ingenuity Knowledge Base |
| ENSMUSG00000037321 | TAP1 | 1.662 | Upregulates | Ingenuity Knowledge Base |
| ENSMUSG00000069516 | LYZ | 1.618 | Upregulates | Ingenuity Knowledge Base |
| **ENSMUSG00000028211** | **TP53INP1** | **1.603** | **Upregulates** | **Ingenuity Knowledge Base** |
| ENSMUSG00000037855 | ZNF365 | 1.511 | Regulates | Ingenuity Knowledge Base |
| ENSMUSG00000026712 | MRC1 | 1.5 | Upregulates | Ingenuity Knowledge Base |
| ENSMUSG00000020326 | CCNG1 | 1.421 | Upregulates | Ingenuity Knowledge Base |
| **ENSMUSG00000026817** | **AK1** | **1.396** | **Upregulates** | **Ingenuity Knowledge Base** |
| ENSMUSG00000041801 | PHLDA3 | 1.393 | Upregulates | Ingenuity Knowledge Base |
| ENSMUSG00000002257 | DEF6 | 1.22 | Regulates | Ingenuity Knowledge Base |
| ENSMUSG00000037999 | ARAP2 | 0.948 | Regulates | Ingenuity Knowledge Base |
| ENSMUSG00000031451 | GAS6 | 0.813 | Upregulates | Ingenuity Knowledge Base |
| ENSMUSG00000030093 | WNT7A | 0.723 | Downregulates | Ingenuity Knowledge Base |
| ENSMUSG00000006356 | Crip2 | 0.486 | Upregulates | Ingenuity Knowledge Base |
| ENSMUSG00000013698 | PEA15 | 0.48 | Regulates | Ingenuity Knowledge Base |
| ENSMUSG00000062762 | EI24 | 0.338 | Upregulates | Ingenuity Knowledge Base |
| ENSMUSG00000001870 | LTBP1 | -0.459 | Upregulates | Ingenuity Knowledge Base |
| ENSMUSG00000092035 | PEG10 | -0.716 | Downregulates | Ingenuity Knowledge Base |

Supplemental Table 7: p53 target genes dysregulated in the *Emx1-Cre;Aki2*$^{fl/fl}$ transcriptome. Bolded genes were validated via qPCR
